# SCONCE2: jointly inferring single cell copy number profiles and tumor evolutionary distances

**DOI:** 10.1101/2022.05.12.491742

**Authors:** Sandra Hui, Rasmus Nielsen

## Abstract

Single cell whole genome tumor sequencing can yield novel insights into the evolutionary history of somatic copy number alterations. Existing single cell copy number calling methods do not explicitly model the shared evolutionary process of multiple cells, and generally analyze cells independently. Additionally, existing methods for estimating tumor cell phylogenies using copy number profiles are sensitive to profile estimation errors. We present SCONCE2, a method for jointly calling copy number alterations and estimating pairwise distances for single cell sequencing data. Using simulations, we show that SCONCE2 has higher accuracy in copy number calling and phylogeny estimation than competing methods. We apply SCONCE2 to previously published single cell sequencing data to illustrate the utility of the method.

## Background

Cancer evolution is driven by an accumulation of somatic point mutations and large copy number alterations (CNAs) (1, 2). Single cell sequencing can offer a detailed picture of the process of CNA and mutation accumulation that is lost in bulk sequencing, in particular by estimating the phylogenetic relationship among different cell types. A challenge in such efforts is that single cell sequencing data is typically very noisy due to variable and low sequencing depth (3), making accurate genotyping, copy number (CN) calling, and phylogeny estimation difficult. However, as we will show here, by leveraging the shared evolutionary history among cells, jointly calling CNAs across cells can lead to increased accuracy and give information about the evolutionary relationship between cells, thereby leading to improved estimates of tumor phylogenies. Different cells from the same tumor share some of their somatic evolutionary history, and information regarding CNAs, and CNA breakpoints, from one cell can, therefore, inform CNA calling in other cells.

Unfortunately, the commonly used methods for estimating single cell copy number profiles (CNPs), the collection of copy number states across the genome, do not rigorously use this shared information. Instead, most methods, including SCONCE (4) and the commonly-used AneuFinder (5, 6), independently call CNPs. Although SCONCE (4), a copy number calling method for single cell tumor data, was previously shown to outperform competing methods in absolute copy number and breakpoint detection accuracy (4), it does not utilize any information from shared evolutionary histories between cells. Other methods, such as CopyNumber (7) and SCIcONE (8), jointly call CNPs by forcing breakpoints to be shared across all cells. However, we showed in previous work that SCONCE (4) outperforms these methods as well, despite not analyzing cells jointly.

Despite these limitations in copy number calling, several distance metrics for copy number profiles have been developed for estimating tumor phylogenies using algorithms such as neighbor-joining (9, 10). Commonly used pairwise distance metrics include the Euclidean distance (11, 12), the MEDICC distance described by (13), and the cnp2cnp distance presented by (14). Although the Euclidean distance is easy to calculate, large and/or overlapping CNAs can artificially inflate this measure, leading to overestimation of dissimilarity. The latter two methods measure distance between two CNPs by attempting to find the minimum number of deletion and amplification events needed to transform one CNP into the other, without allowing regions that are lost to be regained. The MEDICC model is limited to maximum copy number 4 and events that increase or decrease copy number by one, while the cnp2cnp metric relaxes both of these constraints. Cordonnier *et al*. (14) re-implemented the MEDICC algorithm to allow copy numbers greater than 4, and showed that while both the cnp2cnp and MEDICC distances outperform the Euclidean distance for the purpose of phylogeny estimation, cnp2cnp is more accurate on error free data and MEDICC is more accurate on data with errors.

However, none of these methods use explicit evolutionary models of CNAs to provide joint estimates of CNPs and evolutionary distance. Here, we present SCONCE2, an expansion on SCONCE, that further develops SCONCE’s underlying tumor evolutionary model to jointly model the CNA process in two cells. SCONCE2 takes advantage of the shared evolutionary history between cells, and produces more accurate single cell CNP estimates and pairwise estimates of the evolutionary distances between cells, by combining information across multiple cells. We show that SCONCE2 estimates more accurate CNPs and tumor phylogenies than competing methods using extensive simulations, and apply it to previously published data from (11, 15) to illustrate its utility.

## Results

To infer the evolutionary history of tumor cells, SCONCE2 models the evolution of pairs of cells. We assume a pair of cells, (*A, B*), have a partially shared evolutionary history originating from a healthy ancestral diploid cell, *D*. The shared part of their evolutionary history is represented in a tree, 𝒯 = [*t*_1_, *t*_2_, *t*_3_], by a branch of length *t*_1_, running from an non-tumor diploid cell (*D*) to an unobserved divergence point, *Z*. From *Z*, cells *A* and *B* evolve independently, with branch lengths *t*_2_ and *t*_3_, respectively (see Figure 1A). A core goal is to estimate this tree and to distinguish between shared evolutionary events and independent cell specific events.

**Figure 1:**
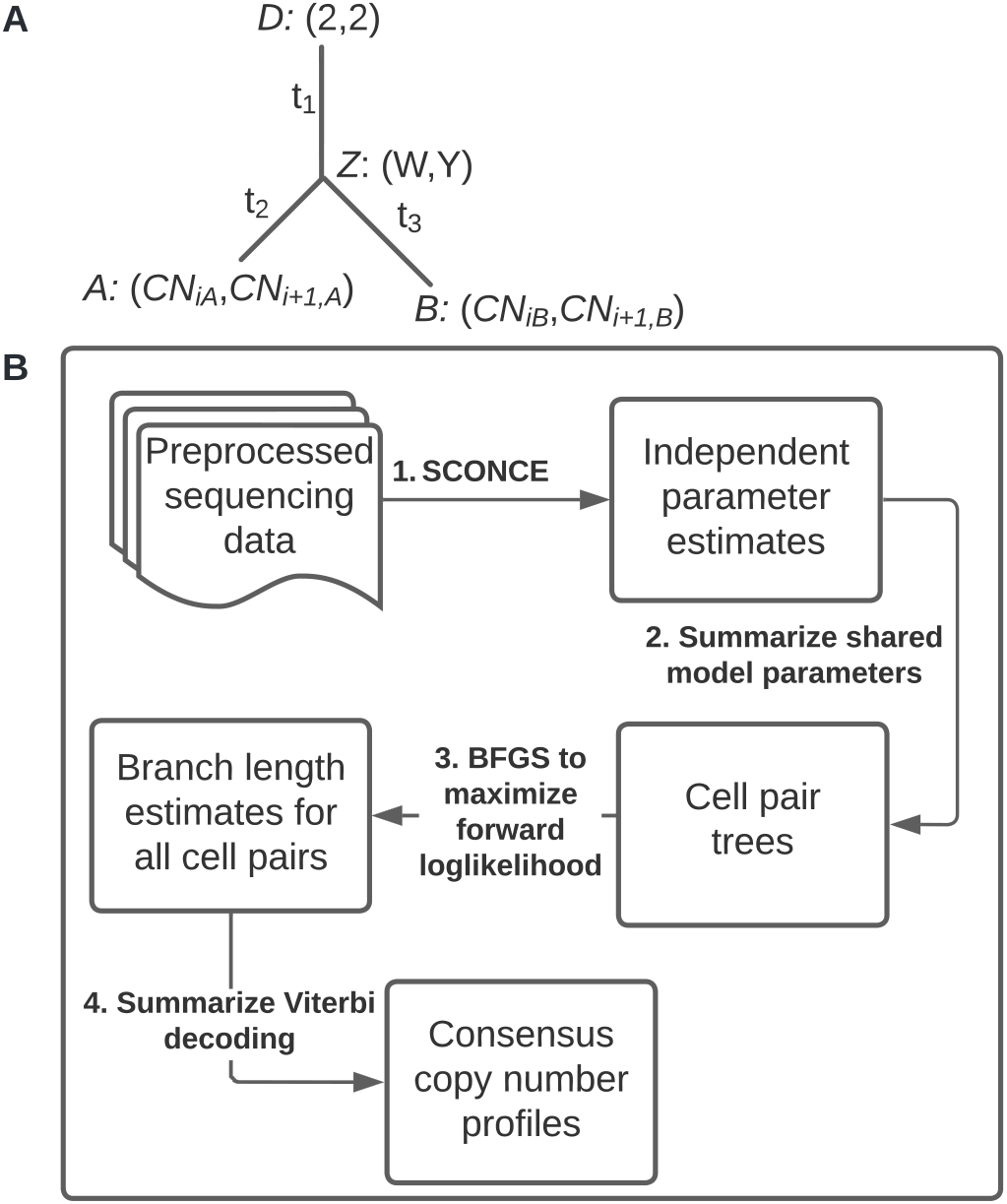
Pairwise tree structure and flowchart of the SCONCE2 workflow. Panel A shows the tree, 𝒯= [*t*_1_, *t*_2_, *t*_3_], between the pair of cells *A* and *B*, where the branch with length *t*_1_ represents their shared evolutionary history from an ancestral diploid cell, *D*, before diverging at the unobserved state *Z*. The branches with lengths *t*_2_ and *t*_3_ show independent evolution to cells *A* and *B*, respectively. Copy number in adjacent bins along the genome is shown in parentheses. Panel B shows the high level workflow of SCONCE2. Each tumor cell is initially independently analyzed through SCONCE, which gives parameter estimates and copy number profiles for each cell. These parameter estimates are then summarized, and branch lengths for tree 𝒯 = [*t*_1_, *t*_2_, *t*_3_] are estimated for each pair of cells. Finally, for each cell, paired copy number profiles are summarized into a consensus copy number profile. Full pipeline details are given in Supplementary Figure S8, Additional File 1.

Because the number of pairs of cells grows quadratically with the number of cells, *n*, full joint maximum likelihood estimation of all parameters can become computationally challenging. We, therefore, first run SCONCE on all cells independently to obtain cell specific estimates of model parameters, and take the median of shared evolutionary parameter estimates across all cells. Then, for each pair of cells, we estimate branch lengths of tree 𝒯 = [*t*_1_, *t*_2_, *t*_3_] using maximum likelihood, and use the Viterbi algorithm to calculate paired decoded copy number profiles. Because each cell appears in *n* − 1 pairs, this produces *n* − 1 paired CNP estimates per cell. Finally, for each cell, we take the per window mean across each cell’s *n* − 1 paired CNP estimates to calculate consensus CNPs. This pipeline is briefly summarized in Figure 1B, with further details given in Methods and Supplementary Figure S8, Additional File 1.

By analyzing each cell in the context of multiple pairs, we obtain increased accuracy in copy number accuracy and breakpoint detection, as well as usable tree branch length estimates. We examine the properties of these estimates results on both simulated and real data.

### Simulations

In order to rigorously test SCONCE2, we applied it to four simulated datasets and two real datasets, from (11, 15). We simulated cells on four different tree structures: tree A) is maximally imbalanced and ultrametric, tree B) is perfectly balanced and ultrametric, tree C) is maximally imbalanced and not ultrametric, where internal and terminal branches have uniform length, and tree D) is maximally imbalanced and not ultrametric, where internal branches have equal length and terminal branch lengths decay logarithmically (tree structures shown in Supplementary Figure S1, Additional File 1). Simulated cells from each tree structure were divided into five discrete subsets of 20 cells each.

Briefly, the simulated genome is modeled as a collection of line segments, where amplifications and deletions occur according to a Markov process and have lengths sampled from a truncated exponential distribution. Copy number events occur within the tree structure, such that ancestral CNAs are propagated to descendent cells. Note, the simulation model is more biologically realistic and intentionally structured to be substantially different from the SCONCE2 inference model, in order to avoid biasing accuracy results to favor our method. We previously described this simulation model in (4), and full simulation details are given in Simulations.

### Copy Number and Breakpoint Detection Accuracy

#### Sum of Squared Error on CNPs

To measure copy number accuracy, we calculated the sum of squared errors (SSE) between the inferred copy number and the true simulated copy number across genomic windows for each cell. To evaluate each step in the SCONCE2 pipeline, we calculated the SSE on copy number profiles generated from individual cell estimation (SCONCE), on profiles from each pair of cells (one pair), and on consensus profiles estimated using three different summary statistics (mean, median, mode). We also compared to AneuFinder (5, 6), a commonly used method for single cell copy number calling, and the second-most accurate one, after SCONCE, among methods evaluated in previous work (4). In all subsequent results, we report summary statistics across all subsets for each tree/simulation set. Recall full simulation descriptions are given in Simulations.

In tree A (maximally imbalanced ultrametric tree; Figure 2A), using pairs of cells had lower SSE than individual cells (SCONCE) alone, with respective median SSE values of 26.01 and 37.31. Furthermore, using the mean had the lowest median SSE of 17.83, with median and mode at 23.68 and 24.04. These SSE values were lower than AneuFinder, which had a median SSE value of 51.78. Similar results for tree B (perfectly balanced ultrametric tree) are shown in Figure 2B, with median SSE values of 28.37, 21.25, 14.74, 17.90, 18.09, and 41.80 for individual cells, single pairs, mean, median, mode, and AneuFinder, respectively. In the same order, the median SSE values for tree C (maximally imbalanced with uniform internal branch lengths) were 73.19, 46.50, 22.20, 32.69, 33.22, and 109.31, and the median SSE values for tree D (maximally imbalanced with logarithmically decaying branch lengths) were 60.13, 40.36, 23.05, 32.00, 33.02, and 76.18 (see Figure 2C, D). Clearly, there is a substantial improvement in accuracy by using pairs of cells instead of individual cells, and this improvement in accuracy is larger if multiple pairs are used.

**Figure 2:**
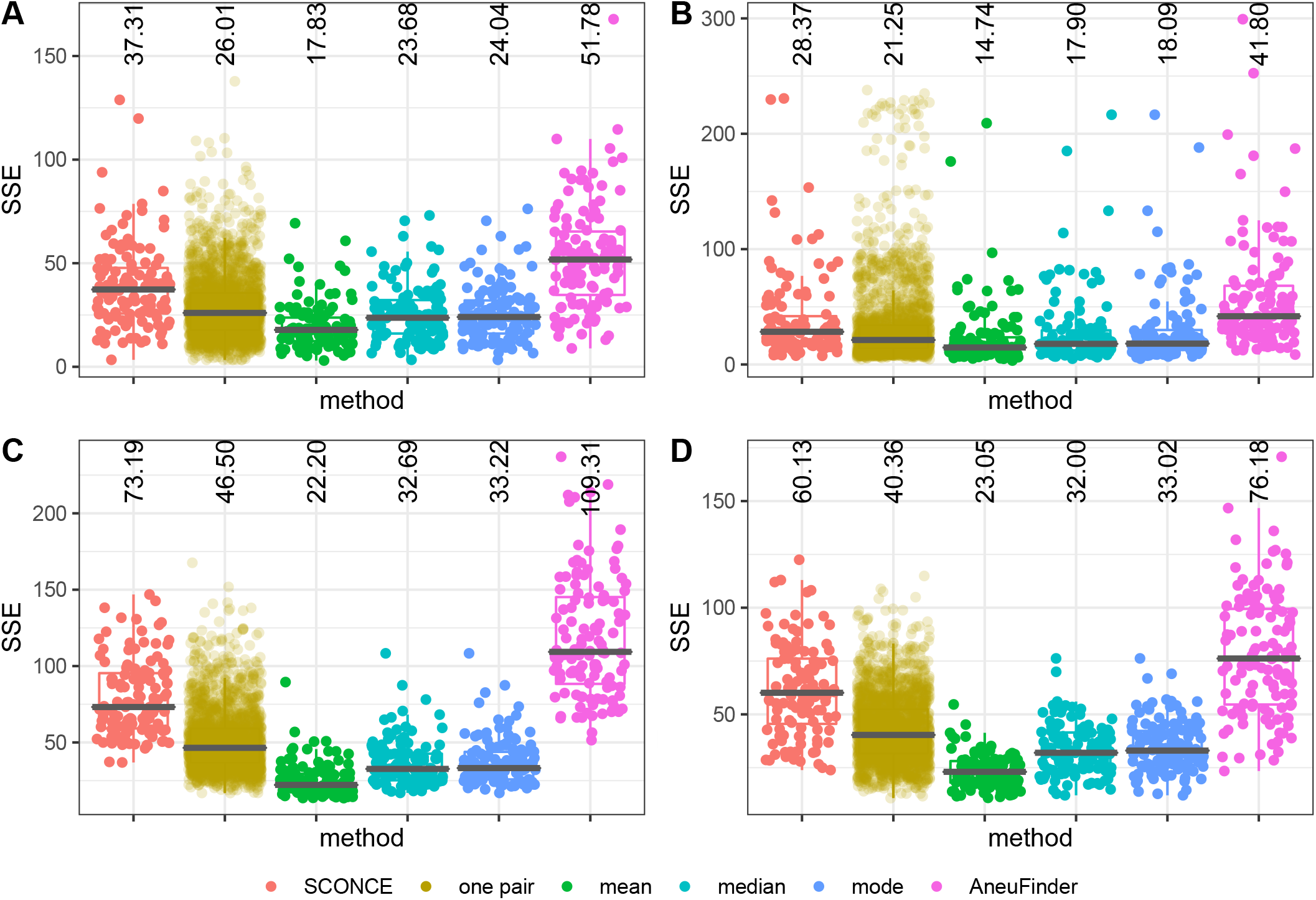
Boxplots of genome wide sum of squared error (SSE) between true simulated copy number profiles and inferred copy number profiles, across methods. SSE results are shown for cell specific CNPs from SCONCE (independent cell inference); joint inference on each pair of cells (one pair); summary functions across all pairs of cells (mean, median, and mode); and AneuFinder. Different methods are shown across the x-axis, and SSE is shown on the y-axis. Median SSE for each method is printed at the top of each column. Each dot represents one cell (note, in “one pair”, each cell appears multiple times), and the median SSE is printed at the top of each column. Panel letters correspond to tree labels A (maximally imbalanced ultrametric tree), B (perfectly balanced ultrametric tree), C (maximally imbalanced non ultrametric tree with uniform branch lengths), and D (maximally imbalanced non ultrametric tree with logarithmically decaying branch lengths). SSE results are consistently lower when using data from multiple cells.

We note that, as an artifact of the genome binning procedure, true fractional copy numbers may occur from small CNAs completely contained within window boundaries, or from CNAs crossing window boundaries (for example, observing windows with true copy numbers 1 → 1.25 → 2). As such, the mean and median have the lowest SSE values because they allow fractional copy numbers. However, many downstream tools expect integer copy number profiles for single cells, so users may wish to round to the nearest integer or use the mode option.

#### Breakpoint distance and detection

In order to measure breakpoint detection accuracy, we calculated the genome wide distance between inferred and true breakpoints, penalized by the number of total inferred breakpoints. Specifically, for each simulated breakpoint, we calculated the distance to the nearest inferred breakpoint. Because erroneously inferring breakpoints at every position in the genome would artificially lower this genome wide distance, we also calculated 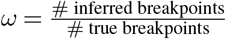, such that lowest breakpoint distances with *ω* values closest to 1 indicate greatest accuracy.

In all simulation sets, using the mean consistently had *ω* values closest to 1, again due to fractional copy number states, as well as lower total breakpoint distance than other methods. Across trees, results from AneuFinder, followed by SCONCE, had the highest breakpoint distances and *ω* values further from 1. For tree A (ultrametric maximally imbalanced tree; Figure 3A), SCONCE, single pairs, mean, median, mode, and Aneufinder had median distance values of 1167, 1006, 394, 1018.5, 1019.5, and 1172, and median *ω* values of 0.466, 0.490, 0.921, 0.486, 0.486, and 0.462, respectively. Similarly, for tree D (maximally imbalanced with logarithmically decaying branch lengths, Figure 3D), median distance values were 153.5, 85, 33, 77, 77.5, and 168.5, and median *ω* values were 0.504, 0.535, 1.007, 0.535, 0.534, and 0.489, in the same order as above. Full median distance and *ω* values are given in Supplementary Tables S2 and S3, Additional File 2. Similarly to the observations for the CNP estimates, breakpoint detection also improves when using pairs of cells, and improves when estimates from multiple pairs are combined, particularly if combining using the mean.

**Figure 3:**
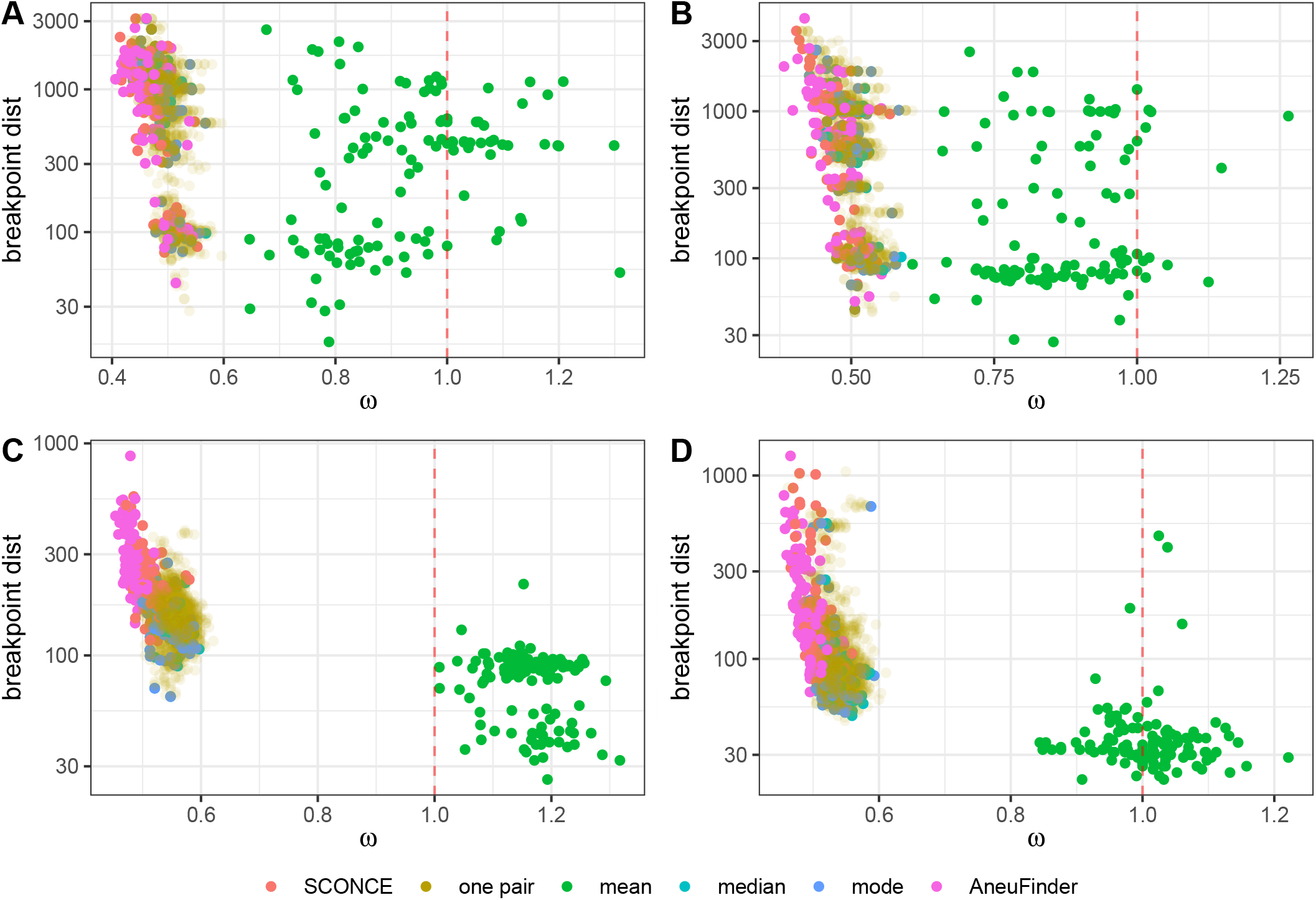
Breakpoint detection accuracy results across methods. Total distance to nearest breakpoint is shown on the y-axis, and 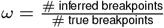 is shown along the x-axis. Each dot represents one cell, colored by method. Methods with the highest breakpoint detection accuracy cluster near *ω* = 1 (vertical red dotted line) and have lowest total breakpoint distance. Each panel corresponds to a simulation set (A-D). In all simulation sets, using the mean, median, and mode have lower total breakpoint distance than independent cell analyses. Furthermore, using the mean results in *ω* values closest to 1, as it is able to infer fractional copy numbers.

#### Optimal number of pairs to use

Because there are 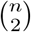 pairs for *n* cells, averaging over more pairs of cells comes at a computational cost. Furthermore, as we will show, adding too many divergent cells can reduce the accuracy, as including highly divergent cells in the average may increase the noise.

To determine the optimal number of cells to summarize across, we estimated summarized (mean) copy number profiles with increasing numbers of cells. As each cell was added, we calculated the difference in SSE relative to SCONCE (individual cells). Cells were added in three different orderings: most to least similar (i.e., nearest first, as defined by the Euclidean distance between the cells’ SCONCE profiles), least to most similar (furthest first), and randomized order. In Figure 4, we summarize the change in SSE across *κ* cells for each tree. Across all trees, SSE improves fastest when adding nearest cells first, and slowest for adding furthest cells first, with the random ordering in between. When adding nearest cells first, the SSE initially sharply decreases, levels off and reaches the largest decrease after approximately 10 cells, and then increases. Specifically, the mean change in SSE when adding nearest cells first reached the greatest decrease in SSE from SCONCE of -20.710, -17.491, -56.185, and -38.570 when *κ* = 12, 10, 9, 15 cells for trees A, B, C, and D, respectively. In contrast, when *κ* = 20 cells, the change in SSE from SCONCE was -19.721, -16.757, -53.461, and -37.920 for trees A, B, C, and D, consistent with the results shown in Figure 2.

**Figure 4:**
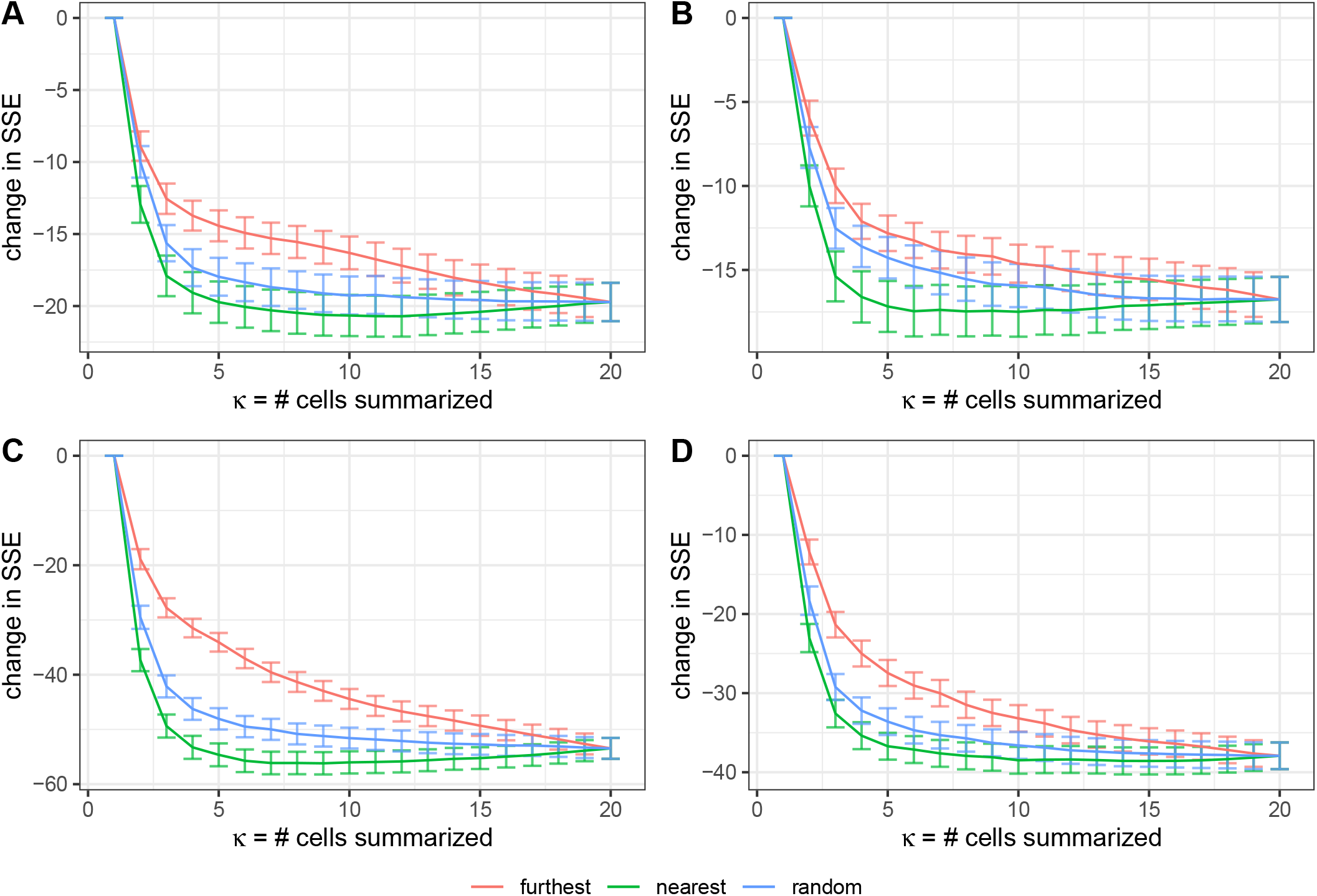
Change in SSE relative to SCONCE, as more cells are added to the consensus (mean) analysis. Number of cells, *κ*, in the joint analysis is shown along the x-axis, and change in SSE (relative to SCONCE) is shown on the y-axis. Each line shows the mean change in SSE across cells, with error bars showing ±1 standard deviation. Colors show cell ordering, where furthest denotes adding the least similar cells first (i.e., furthest distance as defined by the Euclidean distance on SCONCE profiles) and nearest denotes adding the most similar cells first. Across all datasets and cell orderings, SSE quickly drops as more cells are added, with adding the nearest cells (green line) showing the fastest improvement. However, for this ordering, the decrease in SSE levels off after approximately 10 cells are added, and then slightly rises as more cells are added, due to rare copy number events getting averaged out.

When summarizing over *κ < n* − 1 cells, for a given cell, some of the other cells will not be used in that cell’s consensus profiles. As a time saving measure, these excluded cell combinations are not analyzed. Therefore, we recommend users summarize over *κ* = 10 cells, added in order of most to least similar.

For completeness, SSE and breakpoint detection across parameter sets when summarized over only the nearest 10 cells is shown in Supplementary Figures S2 and S3, Additional File 1, and Supplementary Tables S4 and S5, Additional File 2.

#### Using multiple cells results in better CNA detection

Plotting true simulated copy number profiles against inferred copy number profiles shows why performance improves when using multiple cells. For example, in Figure 5, SCONCE erroneously combined two breakpoints for cell A, while predicting cell B’s breakpoint too far to the left (column labelled SCONCE). However, when analyzed as a pair, a shared breakpoint was inferred (left arrow), and the second breakpoint (right arrow) in cell A was correctly inferred (one pair column). While the shared breakpoint was closer to the true breakpoint, it was not until CNPs are summarized across multiple cells that the breakpoint was called in the correct position. Using the mean results in slightly fuzzier boundaries due to non-integer copy number calls (middle column), which better reflected the true underlying data, while the median and mode (right two columns) result in integer jumps at bin boundaries.

**Figure 5:**
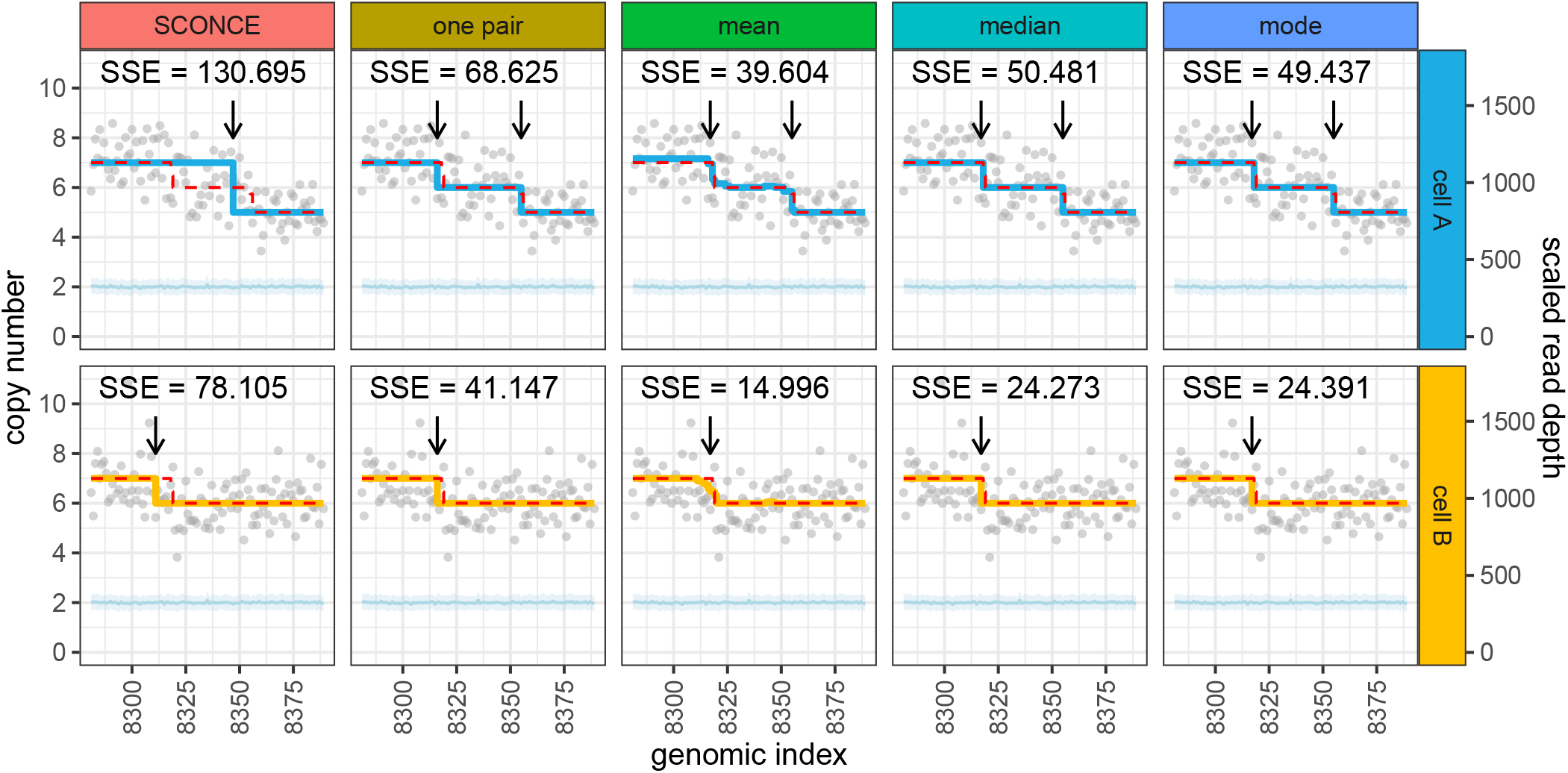
Jointly calling CNAs and summarizing data across multiple cells results in more accurate boundary detection. Inferred copy number profiles and read depth data are shown for two cells (A: 111, B: 59) simulated from tree C (maximally imbalanced, uniform branch lengths). Genomic index shows 110 250kb windows along the x-axis, while the y-axis shows copy number (left) and read depth (right). Gray dots show per window read depth, the light blue line and band show the mean and variance of the diploid read depth, the dotted red line shows the true simulated copy number, and the blue and yellow lines show the inferred copy number calls for each cell. Arrows denote inferred breakpoints, and SSE values are listed for each subpanel. Breakpoint detection accuracy increases as more cells are included in the joint analysis.

Similar results are observed for real data. For example, in Supplementary Figure S4, Additional File 1, SCONCE missed the left most CNA in cell B (arrow in SCONCE column). When analyzed as a pair, this CNA was detected, and was shared with cell A (left arrows). Additionally, a short deletion was called in both cells (right arrows). However, this is a rare event, as it was averaged out in the mean, median, and mode analyses (arrows in mean, median, and mode columns). Furthermore, in Supplementary Figure S5, Additional File 1, SCONCE did not call a CNA in cell A, but did call a -3 deletion in cell B (arrows in SCONCE column). However, when these two cells were jointly analyzed as a pair, there was enough evidence to call a -1 deletion in both. When summarizing across multiple cells, this deletion continued to be supported (right arrow). Additionally, there was some evidence from joint analyses with other cells that an additional small deletion existed in cell B, but not in cell A (left arrow). However, this small deletion was lost when using the median and mode, although the deletion first identified in the joint analysis of cells A and B remained.

### Model Parameter Estimates

For each pair of cells, SCONCE2 estimates the branch lengths for tree 𝒯 = [*t*_1_, *t*_2_, *t*_3_] (see Figure 1B). From the simulated trees, the corresponding tree branch lengths and node distances can be extracted for each cell pair. Recall the simulation and inference models are intentionally formulated differently to evaluate SCONCE2 in more realistic settings (see Simulations). Because the scaling of 𝒯 is different between the simulation and inference models, we show the *R*^2^ values between true (simulated) and inferred values of 𝒯 = [*t*_1_, *t*_2_, *t*_3_], as well as the summed distance *t*_2_ + *t*_3_ as a distance metric between two cells.

For tree A (ultrametric, maximally imbalanced), SCONCE2 recovered {*t*_1_, *t*_2_, *t*_3_, *t*_2_ + *t*_3_} values with *R*^2^ values of 0.798, 0.35, 0.286, and 0.551 (Figure 6A), respectively. Additionally, for tree D (maximally imbalanced with uniform branch lengths), SCONCE2 had *R*^2^ values of 0.661, 0.564, 0.59, and 0.686, for *t*_1_, *t*_2_, *t*_3_, and *t*_2_ + *t*_3_. (Figure 6D).

**Figure 6:**
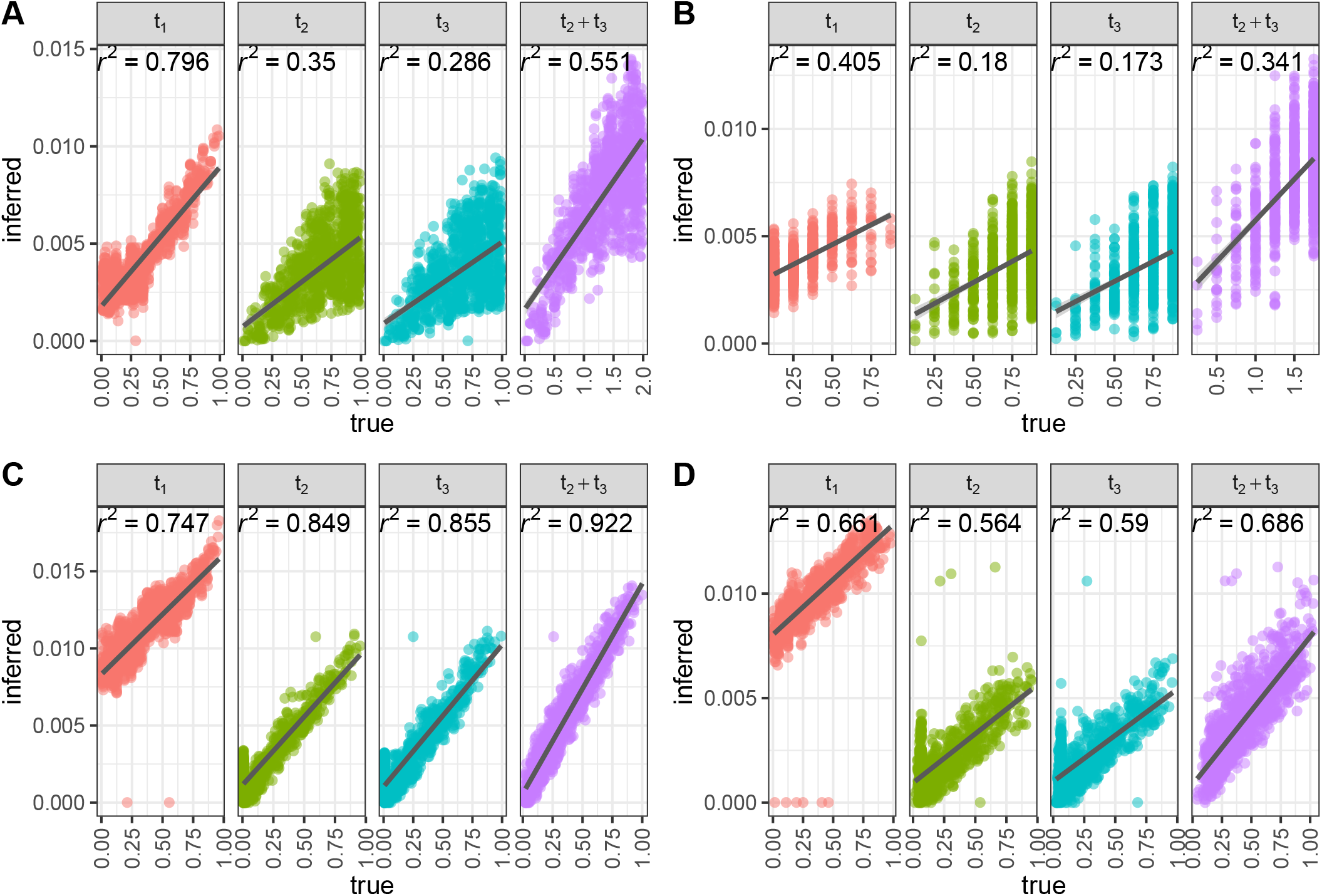
Correlation between true branch lengths and estimated branch lengths across simulation sets. Each dot represents one pairwise branch length estimate, with true node distance on the x-axis and estimated branch length on the y-axis. *R*^2^ values from a linear regression on branch length (dark gray lines) is shown for each subpanel. Across all simulation scenarios (panels A-D correspond to trees A-D), SCONCE2 consistently predicts *t*_1_ and *t*_2_ + *t*_3_ with high *R*^2^ values.

We note that the sum *t*_2_ + *t*3 has higher *R*^2^ values than those of *t*_2_ or *t*_3_ individually, demonstrating some uncertainty in assigning events to particular branches. Furthermore, because the simulation model generates the number of CNAs from a distribution relating to a Poisson (however, the distribution is not truly Poisson as the size of the genome changes through the simulations), the mean and variance of the number of events increases with branch length in expectation. This increased variance is reflected by the larger range of branch length estimates as branch lengths increase. Nonetheless, as we will show in the next section, SCONCE2 recovers the magnitude of cell relationships sufficiently accurately to allow improved phylogeny estimation.

### Phylogeny Estimation

Estimating phylogenies on copy number profiles using neighbor-joining (9, 10) requires a distance metric between cells. Existing metrics include the Euclidean distance (11), and two estimates of the minimum number of CNAs needed to transform one CNP into another: the cnp2cnp metric (14) and the MEDICC distance (13) (here, we use the implementation in the cnp2cnp program (14)). These methods require prior estimation of the CNP. See Running other methods for details on running these programs.

Under the SCONCE2 model, by construction, *t*_2_ + *t*_3_ measures the pairwise distance between two cells. To compare these different distance metrics, we first calculated distance matrices using pairwise Euclidean, cnp2cnp, and MEDICC distances on CNPs called by SCONCE (previously showed to be more accurate than other single cell copy number callers (4)), as well pairwise *t*_2_ + *t*_3_ estimates. Next, we applied neighbor-joining to estimate phylogenies and computed the Robinson-Foulds (RF) distance (16) between the true trees and the inferred trees. As shown in Figure 7, across parameter sets, the trees inferred from estimates of *t*_2_ + *t*_3_ had lower Robinson-Foulds distances than trees inferred from other distance metrics. For example, for tree A (ultrametric, maximally imbalanced), the median RF distances were 27, 27, 29, and 20 for the Euclidean distance, cnp2cnp distance, MEDICC distance, and *t*_2_ + *t*_3_ (Figure 7), respectively (see Supplementary Table S6, Additional File 2, for all median Robinson-Foulds distances).

**Figure 7:**
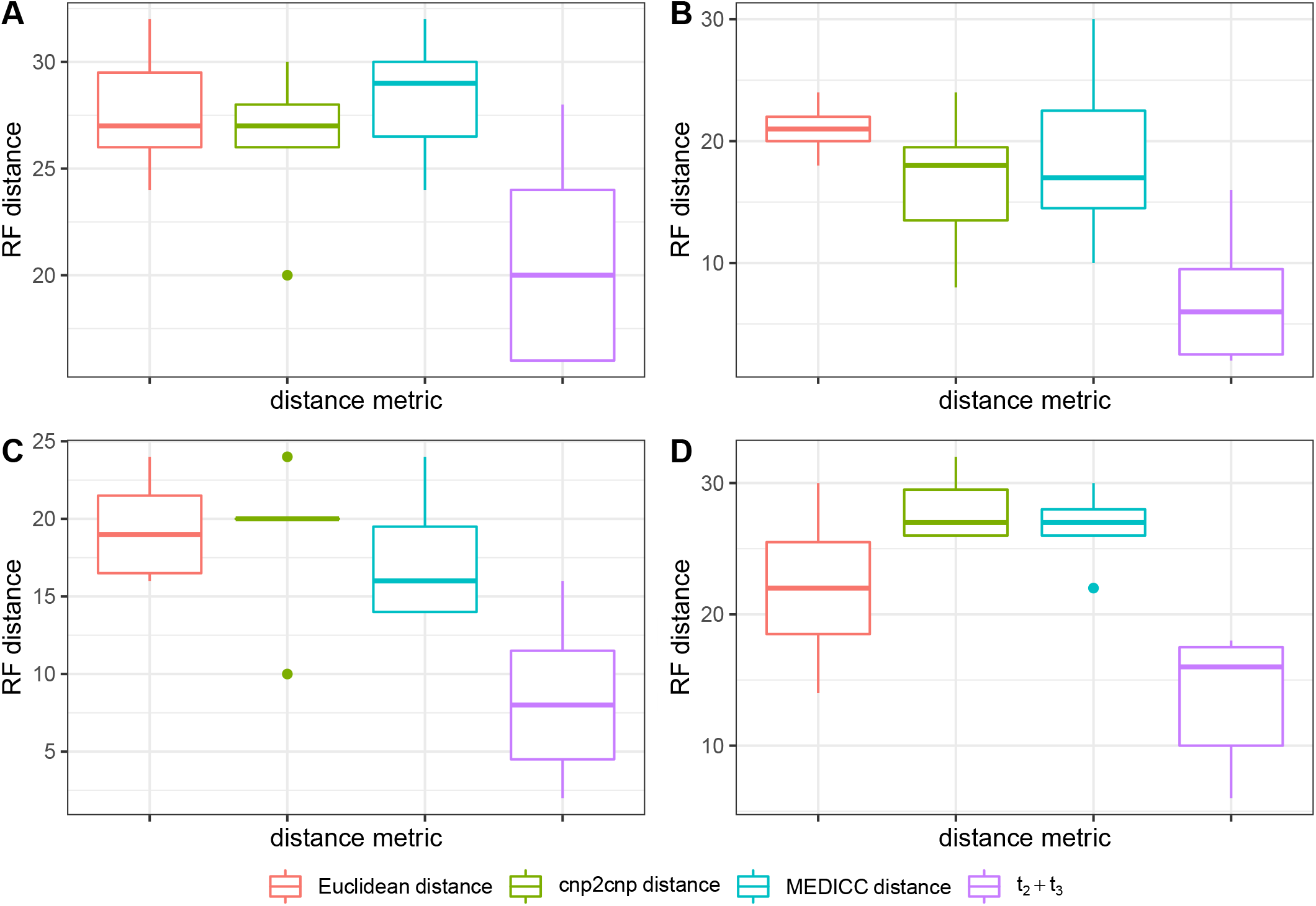
Robinson-Foulds (RF) distances between true and inferred phylogenies. Phylogenies were estimated using neighbor-joining on *t*_2_ + *t*_3_ estimates, and on the Euclidean, cnp2cnp, and MEDICC distances between true CNPs and CNPs inferred from SCONCE. Methods are shown along the x-axis, while Robinson-Foulds distances are shown on the y-axis. Across multiple parameter sets (panels A-D correspond to trees A-D), using *t*_2_ + *t*_3_ estimates resulted in a lower Robinson-Foulds distance from the simulated tree, relative to all other inferred phylogenies.

Furthermore, we calculated RF distances from phylogenies based on the Euclidean, cnp2cnp, and MEDICC distances on consensus CNPs and true simulated CNPs (Supplementary Figure S6, Additional File 1). When summarizing over all pairs of cells, using *t*_2_ + *t*_3_ consistently had lower median RF distances than other methods on consensus CNPs. For example, in tree A (ultrametric, maximally imbalanced), the median RF distances for phylogenies estimated from mean consensus profiles were 27, 26, and 28 for the Euclidean, cnp2cnp, and MEDICC distances (Supplementary Figure S6A, Additional File 1). On distances calculated from the true CNPs, *t*_2_ + *t*_3_ performed as well or better than the other metrics, with the exception of the cnp2cnp distance in tree A, where the Euclidean, cnp2cnp, MEDICC distances and *t*_2_ + *t*_3_ had respective median RF distances of 27, 19, 20, and 20 (Supplementary Figure S6A, Additional File 1). However, under experimental conditions, the true CNP would be unknown. In all other simulation sets, the phylogenies estimated using *t*_2_ + *t*_3_ had lower median Robinson-Foulds distances than all other methods.

For completeness, we additionally calculated Robinson-Foulds distances on phylogenies estimated from consensus CNPs from summarizing over the nearest 10 cells. For tree A (ultrametric, maximally imbalanced), the median RF distances on mean consensus CNPs over the nearest 10 cells were 27, 27, and 26 for the Euclidean, cnp2cnp, and MEDICC distances, respectively (Supplementary Figure S7A, Additional File 1). Note that in order to estimate phylogenies using our *t*_2_ + *t*_3_ metric, all pairs of cells must be analyzed, and cannot benefit from the time savings of analyzing a selected subset of pairs of cells (see Optimal number of pairs to use). This is a weakness of our method, as analyzing all pairs of cells comes at an increased computational cost.

## Discussion

We present a novel method, SCONCE2, that combines data across single cells in a manner that is grounded in a principled model of stochastic tumor evolution. It jointly calls copy number alterations in single cell sequencing of cancer cells with higher accuracy than competing methods on both simulated and real data. Additionally, SCONCE2 calculates an informative pairwise distance metric that can be used to estimate phylogenies with less error than other methods.

Similar to SCONCE, one weakness of SCONCE2 is the requirement for matched diploid cells in order to normalize GC content and mappability biases. These diploid cells must be sequenced under the same experimental conditions for proper normalization and may not be directly of interest to investigators, thereby potentially increasing cost. However, diploid cells are often sequenced as a byproduct of single cell sequencing, and can be identified with orthogonal methods, such as cell sorting.

Additionally, SCONCE2 does not use SNPs or genotype likelihoods, or do any allelic phasing, to inform copy number calls or *t*_2_ + *t*_3_ estimates. Although calling SNPs in low coverage and noisy single cell data is difficult, incorporating genotype likelihoods can add information and increase confidence in these procedures. For example, using the allele frequency in variable sites can support concordant or rule out discordant copy number states, and is the subject of future work.

Another weakness of SCONCE2 is that it takes longer to run, relative to other methods. However, if investigators are primarily interested in copy number calling, significant time can be saved by summarizing over a selective subset of pairs of cells (that is, noninformative pairs are not analyzed), described in Optimal number of pairs to use. But, if investigators are interested in estimating phylogenies using our *t*_2_ + *t*_3_ metric, all pairs of cells must be run in order to create a complete distance matrix (described in Phylogeny Estimation), thereby negating this time saving measure. Nonetheless, we propose the increased accuracy of both copy number calls and phylogeny estimation outweighs the increased computational run time cost.

## Conclusions

In conclusion, we present a principled method, SCONCE2, for simultaneously and accurately calling and aggregating copy number profiles across multiple tumor cells, and estimating pairwise evolutionary distances, using single cell whole genome sequencing. This work shows jointly analyzing cells in single cell experiments to leverage their shared evolutionary history increases accuracy in copy number calling and phylogeny estimation, with implications for deepening our understanding of tumor evolution.

## Methods

### Evolutionary process modeling

We first review the Markov processes introduced in (4). Briefly, we assume an evolutionary process that is continuous in time but discrete along the length of the genome. However, notice that this is just an approximation, as the true process along the length of the genome is not Markovian (see Simulations).

#### One cell continuous time Markov process

The one cell continuous time process from (4) models the copy numbers of two adjacent genomic bins, in positions *i* and *i* + 1 in the genome, on the same lineage (cell) with copy number *U, V* ∈ 𝕊_*c*_ = {0, 1, …, *k*}, respectively, where *k* is the maximum allowed copy number. We assume that (*U, V*) evolve through time with the following rate parameters:

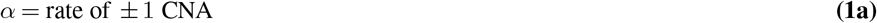

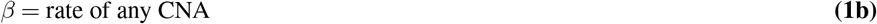

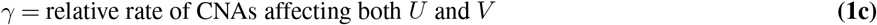

which leads to the following instantaneous rate matrix for the joint process for two bins on one lineage: ℚ = {*q*_(*U,V*),(*U*_′_,*V*_′_)_}:

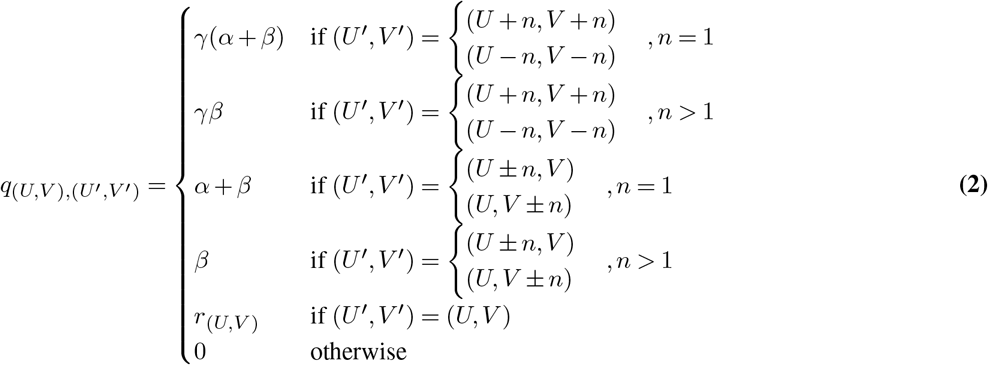

To ensure all rows sum to 0, we set the diagonal elements to the negative row sum, *r*_(*U,V*)_ = −Σ_(*u*′,*v*′) ≠ (*U,V*) *q*(*U,V*),(*u*′,*v*′)_. Note that ℚ defines the instantaneous rate of events such as (*U*′, *V*′) = (*U* + *n, V* − *k*), *n >* 0, *k >* 0 (i.e., events where (*U, V*) are changed by different CNAs) to be equal to 0. However, (*U, V*) can be changed by different CNAs for any evolutionary time interval *t >* 0.

The corresponding probability matrix, ℙ, of time dependent transition probabilities of adjacent bins changes from (*U, V*) to (*U*′, *V*′) is calculated as the matrix exponential

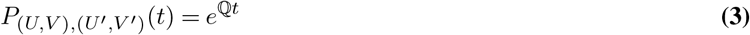

where *t* is evolutionary time.

#### Two cell evolutionary process expansion

We now extend this single lineage process to describe the joint evolutionary process in two cells. Consider a pair of cells (*A, B*) and their most recent common ancestor in a tree, 𝒯 = [*t*_1_, *t*_2_, *t*_3_], where *t*_1_ denotes the branch length of their shared history and *t*_2_ and *t*_3_ denote the branch lengths from divergence, at unobserved state *Z*, to cells *A* and *B*, respectively (see Figure 1A).

Under this tree structure, adjacent bins in cells *A* and *B* have a shared evolutionary history for time *t*_1_ from an ancestral diploid state (i.e., *D* : (2, 2)) to an intermediate unobserved state, *Z* : (*W, Y*), with associated transition probability *P*_(2,2),(*W,Y*)_(*t*_1_). After divergence, bins in cell *A* evolve from (*W, Y*) to (*CN*_*iA*_, *CN*_*i*+1,*A*_) in time *t*_2_ with transition probability *P*_(*W,Y*),(*CNiA*_,*CN*_*i*+1,*A*_)(*t*_2_), where *CN*_*iA*_, *CN*_*i*+1,*A*_ ∈ 𝕊_*c*_ denote copy number in windows *i* and *i* + 1 for cell *A*. Similarly, bins in *B* evolve from (*W, Y*) to (*CN*_*iB*_, *CN*_*i*+1,*B*_) in time *t*_3_ with transition probability (*W, Y*), (*CN*_*iB*_, *CN*_*i*+1,*B*_) (*t*_3_).

#### Approximating discrete Markov process along the genome

Next, we convert these continuous time process transition probabilities for adjacent bins in two cells into the transition probabilities for the approximating discrete Markov process for pairs of cells along the entire length of the genome, further described in Two Cell Hidden Markov Model Description. We do this by expanding the state space to the product space of the state space for each cell, 𝕊_*c*_. This expansion of the state space to the joint CN state for two cells is necessary as the correlation structure along the length of the genome prevents the use of standard tree-based dynamic programming algorithms such as Felsenstein’s pruning algorithm (17).

The state space, 𝕊_*d*_, for this discrete process is composed of pairs (*CN*_*iA*_, *CN*_*iB*_), representing the copy numbers for window *i* in cell pair (*A, B*), where *CN*_*iA*_, *CN*_*iB*_ ∈ 𝕊_*c*_ = {0, 1, …, *k*} for a fixed maximal copy number *k*, such that

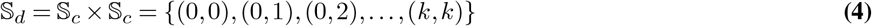

We define matrix 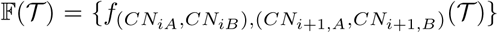 as the transition probability of moving from state (*CN*_*iA*_, *CN*_*iB*_) in window *i* to (*CN*_*i*+1,*A*_, *CN*_*i*+1,*B*_) in window *i* + 1, given evolutionary tree 𝒯 = [*t*_1_, *t*_2_, *t*_3_], for cell pair (*A, B*). Therefore, the matrix 𝔽(𝒯) is defined as

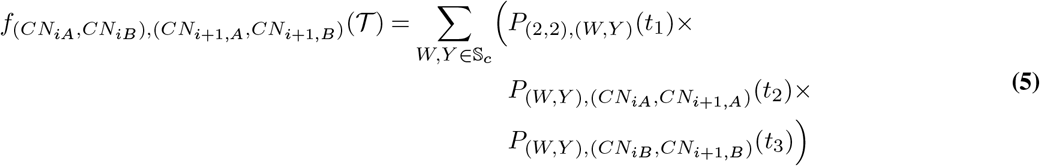

which can be used to calculate a transition matrix, 𝕄(𝒯), along the length of the genome for pairs of cells. This is done by dividing the joint probability of the CN state in both cells (*A* and *B*) in both bins (*i* and *i* + 1), with the marginal probability of CN state in both cells (*A* and *B*) in bin *i*, i.e., dividing each entry in 𝔽(𝒯) with the corresponding row sum:

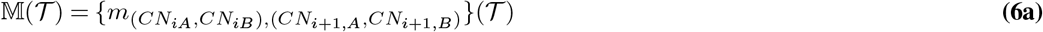

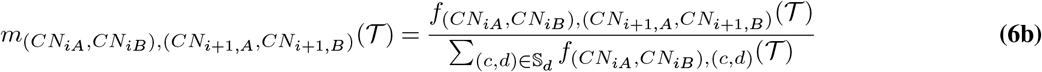

We have thereby constructed a process with state space on the copy numbers of pairs of cells, 𝕊_*d*_. The matrix 𝕄(𝒯) gives the probabilities of observing transitions from (*CN*_*iA*_, *CN*_*iB*_) in window *i* to (*CN*_*i*+1,*A*_, *CN*_*i*+1,*B*_) in window *i* + 1, along the genome, for cell pair (*A, B*), given evolutionary tree 𝒯. We also note that the process along the length of the genome is not Markovian (see also (4)). However, to facilitate computation, we will approximate the process as a Markovian process with transition probabilities given by 𝕄(𝒯). This Markov chain will then be used for inferences in a Hidden Markov Model framework with emission probabilities similar to those described in (4).

### Two Cell Hidden Markov Model Description

Expanding on the framework of (4), we define a Hidden Markov Model (HMM) (18–20) to infer copy number across the genome for pairs of tumor cells, using binned read depth data.

Recall that cells *A* and *B* are associated with the evolutionary tree, 𝒯, shown in Figure 1A. The sample space of read data, 𝔸, is composed of pairs of observed per window read count values, (*x*_*iA*_, *x*_*iB*_), *x*_*iA*_, *x*_*iB*_ ∈ ℕ_0_ = {0, 1, 2, …}:

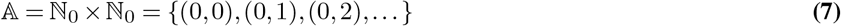

This HMM uses the state space of paired copy numbers, 𝕊_*d*_, defined in Equation 4 and the transition matrix 𝕄(𝒯), defined in Equation 6.

#### Emission probabilities

Assuming conditional independence between cells, the emission probabilities of the HMM are:

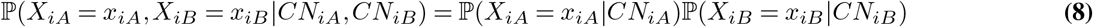

As these probabilities are calculated similarly for cells *A* and *B*, we only describe the derivation for cell *A* (note: we previously described this derivation in (4)).

We assume *X*_*iA*_ follows a negative binomial distribution, such that

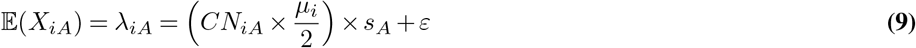

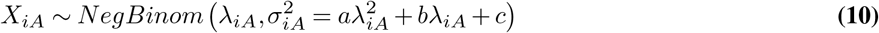

where

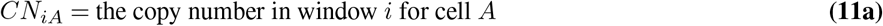

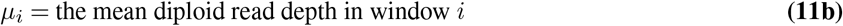

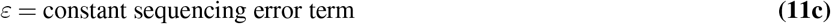

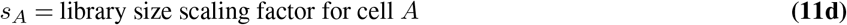

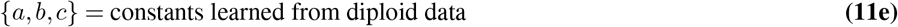

Estimation of constants {*a, b, c*} is described in Initial independent parameter estimation using SCONCE. In the following, we will describe the full SCONCE2 estimation procedure in detail.

### Detailed SCONCE2 pipeline

Given binned read depths for tumor and matched diploid cells, joint copy number calling in SCONCE2 takes place in four main steps: 1) independently estimating model parameters and copy number profiles for each cell using SCONCE, 2) combining independent parameter estimates across cells, 3) estimating tree branch lengths for each cell pair, and 4) creating summarized copy number profiles. This process is illustrated in Supplementary Figure S8, Additional File 1.

#### Initial independent parameter estimation using SCONCE

We first estimate constants {*a, b, c*}, defined in Equations 10 and 11e, using maximum likelihood on diploid cells only, as previously described in (4). We note that most single cell tumor sequencing projects naturally also produce data from non-tumor diploid cells as part of standard sequencing techniques, and that these cells conveniently can be used for standardization (11, 21–27).

In order to obtain initial estimates of all model parameters, we analyze all tumor cells independently through SCONCE, described in detail in (4) and briefly summarized here. This is done to avoid the computational cost of joint estimation for all model parameters across all pairs of cells. The SCONCE pipeline first estimates the transition matrix of an unconstrained CN HMM, with associated library size scaling factor *s*_*A*_, for each cell using a modified Baum-Welch (28) algorithm. These estimates are then used to obtain initial starting points for each model parameter for an optimization of the likelihood function using the Broyden–Fletcher–Goldfarb–Shanno (BFGS) algorithm (29). This results in parameter estimates 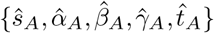, for cell *A*. Recall *s*_*A*_ is the library size scaling factor, defined as the coverage for the cell relative to the average diploid library size, {*α, β, γ*} are the instantaneous rates for copy number events, and *t*_*A*_ is the total branch length from the ancestral diploid cell to cell *A* (see the red block in Supplementary Figure S8, Additional File 1.).

#### Combining parameter estimates across multiple cells

To analyze shared evolutionary history between *n* cells, we first combine independent estimates across all cells of the transition rate parameters, {*α, β, γ*}, assumed to be shared among all cells, using the median to form joint estimates:

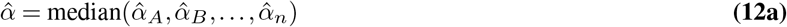

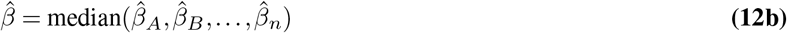

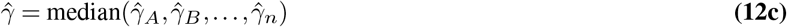

While a full joint optimization could possibly lead to better estimates, we opted not to pursue the estimation of such estimates because of considerations of computational efficiency. This is illustrated in the yellow block in Supplementary Figure S8, Additional File 1.

#### Estimating pairwise tree branch lengths

Next, we estimate parameters of the joint two-cell process for all 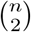 pairs of cells. The branch lengths of tree 𝒯 = [*t*_1_, *t*_2_, *t*_3_], are specific to each pair of cells, and branch length estimates from SCONCE, 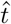, are used to inform the initial optimization starting point for 𝒯. For example, for pair (*A, B*), the initial branch length estimates, denoted with ^*^, are:

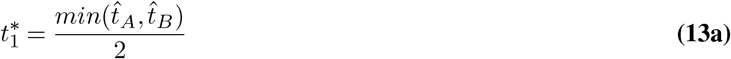

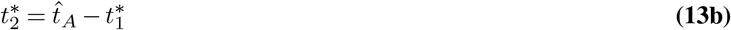

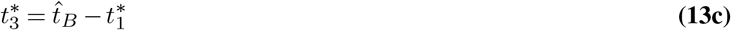

For each pair of cells, we use the BFGS algorithm to maximize the forward log likelihood in order to estimate 𝒯. To calculate the forward log likelihood of an observed sequence, the HMM is reset into the initial probability vector, defined as the steady state distribution, at the beginning of each chromosome to ensure chromosomal independence.

Because each set of branch lengths, [*t*_1_, *t*_2_, *t*_3_], is specific to each pair of cells, this procedure is trivially parallelizable (see the green block in Supplementary Figure S8, Additional File 1).

#### Summarized copy number calling

After pairwise branch lengths are estimated, we use the Viterbi algorithm (30, 31) to estimate the most likely joint copy number profile for each pair of cells. If cell *A* appears in *n* − 1 pairs, this results in *n* − 1 separate CNP estimates for cell *A*. In order to calculate a single consensus copy number profile, *CN*_*A,consensus*_, we use either the mean (default), median, or mode of the CN in each window among the *n* − 1 estimates.

While adding more information to each consensus copy number profile by summarizing across multiple cells initially increases accuracy, summarizing across too many divergent cells is not optimal because more accurate estimates about each cell are obtained using closely related cells than highly divergent cells (Figure 4). Therefore, in order to balance combining data from multiple cells and maintaining cell specificity, the user can also choose to summarize across a subset of the *κ* nearest neighbors for each cell, instead of all *n* − 1 pairs a particular cell appears in. The nearest neighbors for each cell is defined by the Euclidean distance between individual copy number profiles from SCONCE. Then, the consensus copy number profile is calculated only across the *κ* selected pairs.

Note, because of the genomic binning procedure, true copy number events may be split across bin boundaries or be completely contained within one bin, resulting in bins with non-integer average copy number. Using the mean and median summary functions can result in non-integer copy number calls, which more accurately represent the underlying biology as genomes are not truly organized in discrete bins. However, many downstream tools for single cell analyses require integer copy number profiles, so these values may need to be rounded for downstream analyses.

### Simulations

In order to evaluate the accuracy of SCONCE2, we use the Line Segments model from SCONCE (4). However, instead of using coalescent simulations, we specify two ultrametric trees with uniform branch lengths, where tree A is fully pectinate/maximally imbalanced and tree B is perfectly balanced, and two non ultrametric trees, where tree C has uniform internal and terminal branch lengths, and tree D has uniform internal branch lengths and logarithmically decaying terminal branch lengths. For illustration, the tree structure for 8 cells is shown for each dataset in Figure S1. In order to ensure tree B was a perfectly balanced binary tree and to be consistent between tree structures, read depths for 128 tumor cells and 100 diploid cells were simulated for each tree. Read depth across diploid cells was averaged per window for each tree. Tumor cells from each tree were divided into five non overlapping subsets of 20 cells to create test sets. Although healthy cells were shared for each analysis run, each test set was otherwise analyzed independently from other test sets from the same tree.

Total tree heights were all scaled to 1, and simulated genome lengths were set to 100, with amplification and deletion rates and expected lengths shown in Supplementary Table S1, Additional File 2 (relative to the genome length). For read simulation, the human reference genome was divided into 12,397 windows (number of 250kb non overlapping uniform windows in hg19), and read depths were simulated from a negative binomial distribution with parameter *r* = 50 and 4,000,000 total expected number of reads for each cell. All parameter files used to generate simulations are available on GitHub.

### Real data preprocessing

We applied SCONCE2 to two published breast cancer, known for their frequent CNAs (32), single cell datasets, from (11) and (15). Both of these datasets were processed as previously described (4). Briefly, for the (11) dataset, we trimmed reads using cutadapt (33) and trimmomatic (34), removed low complexity reads with prinseq (35), aligned reads to hg19 using bowtie2 (36), removed reads with q scores less than 20 using samtools (37), and removed PCR duplicates using picard (38). For the (15) dataset, we split downloaded preprocessed bam files into cell specific bam files using pysam (39), and removed reads with q scores less than 20 using samtools (37). Finally, we used bedtools to count per window read depth for each cell (40). Cells previously and orthogonally identified as diploid cells in (11) served as the matched normal. For the (15) dataset, subset A was used as the diploid samples, as previous described (26).

### Running other methods

For benchmarking, we limit our comparisons to other copy number only methods (that is, no SNP or phasing information is used): SCONCE (4) and AneuFinder (5, 6) for copy number accuracy, and the cnp2cnp (14) and MEDICC (13) distances for phylogeny building.

Briefly, we ran AneuFinder with default parameters, with the exception of skipping GC and mappability corrections to avoid overcorrecting, as we did not include GC or mappability bias in our simulations. To benchmark SCONCE2’s copy number calling, we first ran SCONCE (4) with default parameters (`k=10`). To run AneuFinder (5, 6), we skipped the GC and mappability corrections steps to avoid over correcting, as our simulation model does not include GC or mappability biases. We directly ran AneuFinder’s `findCNVs` function (default parameters: `method=“edivisive”, R=10, sig.lvl=0.1`). We extracted copy number calls from the resulting the `copy.number` element, and used `bedtools intersect` (40) to split large segments into 250kb windows.

To evaluate SCONCE2’s *t*_2_ + *t*_3_ distance metric in phylogeny estimation, we compared to the cnp2cnp distance (14) and the MEDICC distance (13). To run cnp2cnp, we first converted and rounded called CNPs into fasta files, then ran cnp2cnp in matrix mode with default parameters (`-m matrix -d any`). Because the cnp2cnp metric depends on the input sample ordering and is not symmetric, we repeated this process on the reversed sample ordering, and summed the two resulting distance matrices to make a symmetric metric. To calculate the MEDICC distance, we used the ZZS implementation of the MEDICC algorithm in the cnp2cnp program to remove the maximum copy number limit of 4 in the original MEDICC software, and ran it on the same fasta files (`-m matrix -d zzs`). Full scripts to run other methods are provided on GitHub.

### Phylogeny estimation and Robinson-Foulds distance calculations

To estimate phylogenies, distance matrices were first read into R (41). Next, we applied neighbor-joining (9, 10), implemented in the `ape` (42) package, to each distance matrix to estimate phylogenies.

To calculate Robinson-Foulds distances between inferred trees and true trees, we first used the `read.tree` function in the `ape` (42) package to read true trees in Newick format into R. Next, we used the `treedist` function from the `phangorn` (43, 44) package to calculate the Robinson-Foulds distance. Full scripts to estimate phylogenies from distance matrices and calculate Robinson-Foulds distances between phylogenies are available on GitHub.

## Abbreviations

BFGS: Broyden–Fletcher–Goldfarb–Shanno algorithm (29)
CN: copy number
CNA: copy number alteration(s)
CNP: copy number profile
HMM: hidden Markov model (18–20)
RF distance: Robinson-Foulds distance (16)
SNP: single nucleotide polymorphism
SSE: sum of squared errors

## Appendix

### Ethics approval and consent to participate

Not applicable

### Consent for publication

Not applicable

### Availability of data and materials

SCONCE2 is implemented in C++11, requires the Boost C++ Libraries (developed on v1.65.1)and the GNU Scientific Library (developed on v2.4) (45), has been developed and tested on Ubuntu 18.04.6, and is freely available from https://github.com/NielsenBerkeleyLab/sconce2. The simulation program (written in C) and corresponding parameter files, all R scripts (developed on R v4.1.2) (41) needed to preprocess diploid data (`avgDiploid.R` and `fitMeanVarRlnshp.R`), and R plotting scripts (`readBedFilesPairs.R, readTreeBranches.R, plotBetterBoundaries.R, plotDiminishingReturnsNumPairs.R, plotIllustrativeTrees.R, plotRFdist.R, plotSSEandBreakpointPairs.R, plotTreeBranchCorrelation.R`) are also on GitHub. Plotting scripts require the R packages `ape` (42), `cowplot` (46), `ggplot2` (47), `ggtree` (48–50), `grid` (41), `gtools` (51), `phangorn` (43, 44), plyr (52), `reshape2` (53), `scales` (54), and `stringr` (55).

We analyzed two previously published real datasets. Data from (11) is available at the Sequence Read Archive (SRA) under accession number SRR054616. The 10x dataset from (15) is available at https://cf.10xgenomics.com/samples/cell-dna/1.1.0/breast_tissue_aggr_10k/breast_tissue_aggr_10k_web_summary.html.

### Competing interests

The authors declare that they have no competing interests.

### Funding

This work was supported by the National Institutes of Health [R01GM138634-01 to R.N.].

### Authors’ contributions

SH designed and implemented the SCONCE2 pipeline, and performed all data analysis. RN conceived the study and developed the simulation program. SH and RN wrote the manuscript.

