## Supplementary Notes, Figures, and Tables for "SCONCE2: jointly inferring single cell copy number profiles and tumor evolutionary distances"

### Supplementary Notes and Figures

**Tree illustrations.** We simulated 128 cells from four distinct tree structures, described in Simulations. Small 8 cell trees are shown for illustration purposes in Supplementary Figure S1, Additional File 1. Tree A is ultrametric and maximally imbalanced, tree B is ultrametric and perfectly balanced, tree C is not ultrametric (all branches have uniform length 1/128) and maximally imbalanced, and tree D is not ultrametric (all internal branches have uniform length 1/128, and all terminal branches have logarithmically decaying branch lengths) and maximally imbalanced.

**SSE and breakpoint distance accuracy over the nearest 10 cells.** Across parameter sets, the median sum of squared error (SSE) between the true and inferred copy number profile for each cell is lowest when each cell's consensus profile is calculated by summarizing across the nearest 10 cells, rather than over all 20 in each subset, as rare events aren't averaged out. The nearest cells are determined by the lowest Eucliden distance on each cell's SCONCE profile. Similar to Figure 2, SSE is shown across parameter sets for CNPs from SCONCE (independent inference); from each pairwise analysis; summarized across pairs using mean, median, and mode; and from AneuFinder in Supplementary Figure S2, Additional File 1.

Additionally, similar to Figure 3, we show breakpoint detection accuracy in Supplementary Figure S3, Additional File 1 when consensus profiles are calculated by summarizing over each cell's nearest 10 cells. Median breakpoint distance and $\omega = \frac{\#\text{ inferred breakpoints}}{\#\text{ true breakpoints}}$ values are given in Supplementary Tables S4 and S5, Additional File 2.

**Median Breakpoint Distance and $\omega$ Values.** To evaluate breakpoint detection accuracy, we summed the distance from each true breakpoint to its nearest inferred breakpoint. We define $\omega = \frac{\#\text{ inferred breakpoints}}{\#\text{ true breakpoints}}$, such that highest accuracy is indicated by low breakpoint distance and $\omega$ close to 1. Median breakpoint distance and $\omega$ values are given across all analyzed cells for each tree in Supplementary Tables S2 and S3, Additional File 2. Similarly, median breakpoint distance and $\omega$ values when consensus methods only summarize across the 10 nearest cells are given in Supplementary Tables S4 and S5, Additional File 2.

**Robinson-Foulds Distance Values.** SCONCE2 estimates branch lengths for tree $\mathcal{T} = [t_1, t_2, t_3]$ for every pair of cells. To evaluate the usefulness of $t_2 + t_3$ as a distance metric for phylogeny building, we first calculated distance matrices using $t_2 + t_3$. For comparison, we calculated distance matrices using several different distance metrics (Euclidean distance, cnp2cnp, and MEDICC) at different points in the SCONCE2 pipeline (after SCONCE (independent inference); on summarized CNPs from the mean, median, and mode). As a sanity check, we also calculated distance matrices on the true simulated CNPs (i.e., removing all noise from the copy number inference process). Next, we applied neighbor-joining to these distance matrices and calculated the Robinson-Foulds (RF) distance between each resulting tree and the true simulated tree. For each cell subset in each tree, RF distances are plotted in Supplementary Figure S6, Additional File 1, and median RF distances are given in Supplementary Table S6, Additional File 2.

Using the Euclidean distance is remarkably stable to variations in CNPs. Both the cnp2cnp and MEDICC metrics had the highest median RF distances for SCONCE, and the lowest on the true CNPs, with distances from mean, median, and mode lying between those two extremes. Across all parameter sets, using $t_2 + t_3$ resulted in median RF distances lower than other methods, with the exceptions of using the true CNPs. However, it stands to reason that one would not have access to noiseless copy number profiles in experimental conditions.

Additionally, we calculated the above distance matrices and ran neighbor-joining on CNPs where consensus profiles were summarized across only the nearest 10 cells. However, analyzing only the nearest 10 cells means some cell pairs get skipped, and therefore do not have an $t_2 + t_3$ estimate. To address this, we only calculated RF distances on trees estimated from other metrics.RF distances are plotted in Figure S7, and median RF distances are given in Supplementary Table S7, Additional File 2.

**Detailed SCONCE2 pipeline.** The SCONCE2 pipeline, shown in Supplementary Figure S8, Additional File 1, consists of four main steps: 1) Tumor cells are first independently analyzed through SCONCE to estimate model parameters (for cell $A$, $\{s_A, \alpha_A, \beta_A, \gamma_A, t_A\}$) and cell specific copy number profiles. 2) Shared parameter estimates, $\{\alpha, \beta, \gamma\}$, are then summarized across all cells using the median. 3) The branch lengths in tree $\mathcal{T} = [t_1, t_2, t_3]$ are then independently estimated for each pair of cells. Finally, the Viterbi decoding is used to calculate copy number profiles for each pair of cells, and 4) cell specific copy number profiles are summarized across pairs. Optionally, only the $\kappa$ nearest neighbor cells (defined by the Euclidean distance on the SCONCE copy number profiles) are summarized for a given cell.

DRAFT

### Supplementary Figures

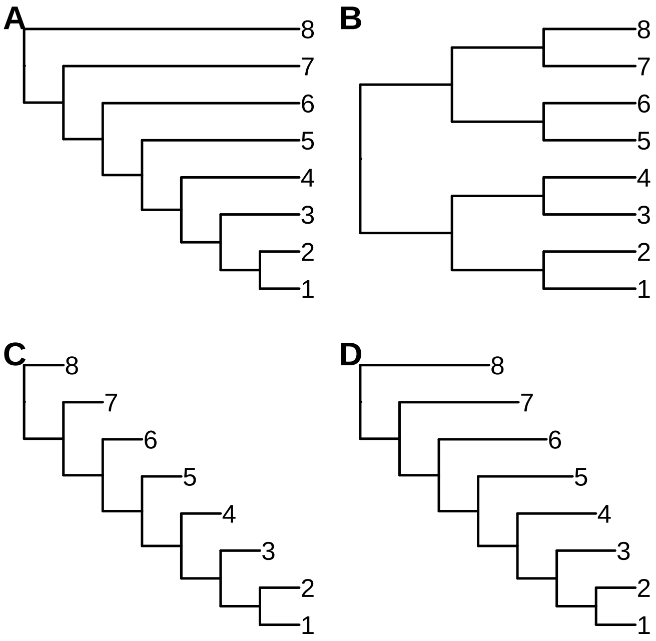


Figure S1: Tree structure for each simulated data set. Although each tree contained 128 cells, only 8 cells for each tree is shown for brevity.

DRAFT

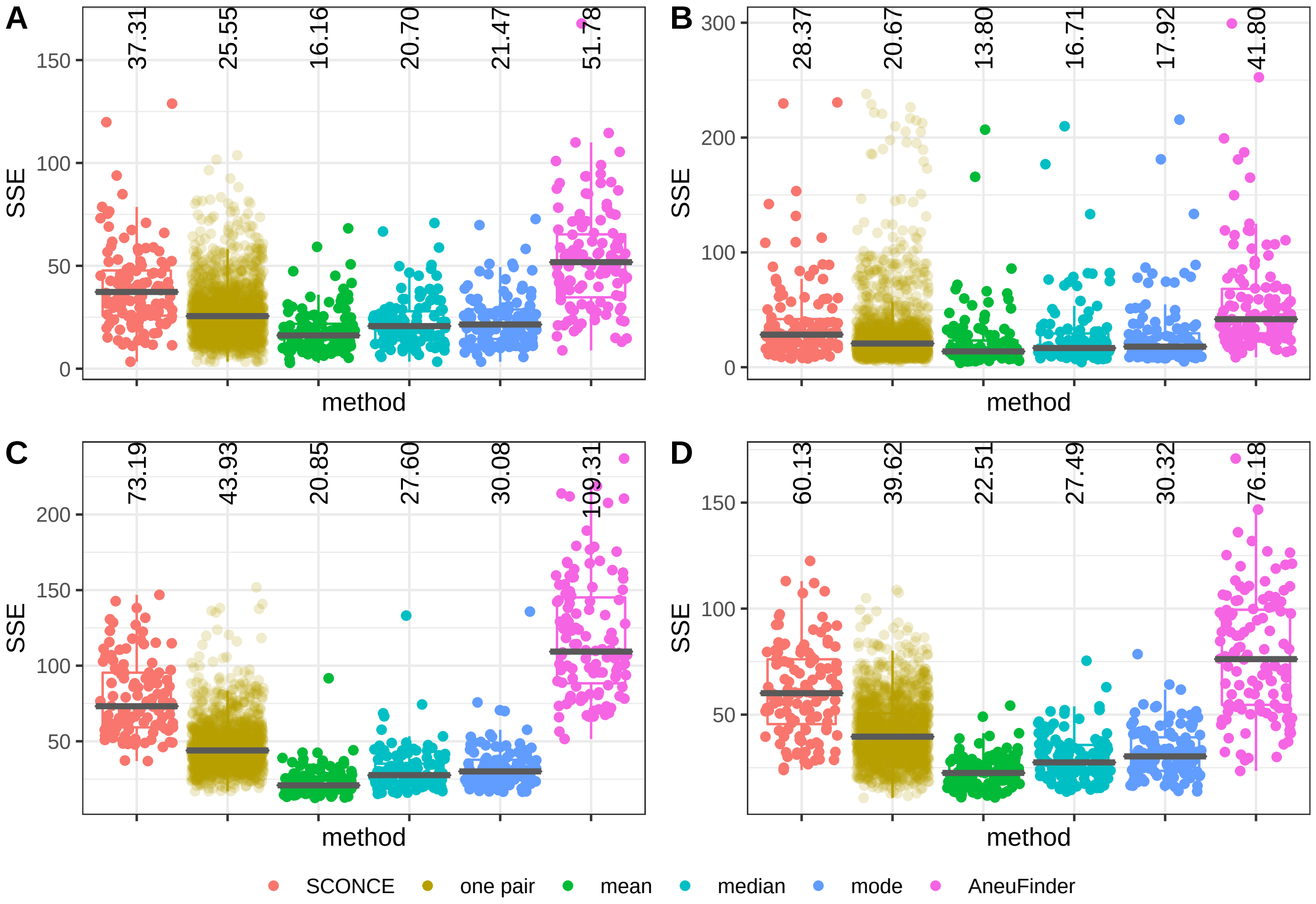


Figure S2: Boxplots of SSE across parameter sets and CNP calling method. Here, for summary methods (mean, median, and mode), each cell's consensus CNP is calculated over its nearest 10 cells. Each dot shows one cell, CNP calling method is on the x-axis, and SSE is on the y-axis, with median SSE values printed at the top of each column. The median SSE is indicated with a black bar and printed at the top of each column. Across parameter sets, calculating consensus calls results in lower SSE than both competing methods and summarizing over all pairs of cells (Figure 2).

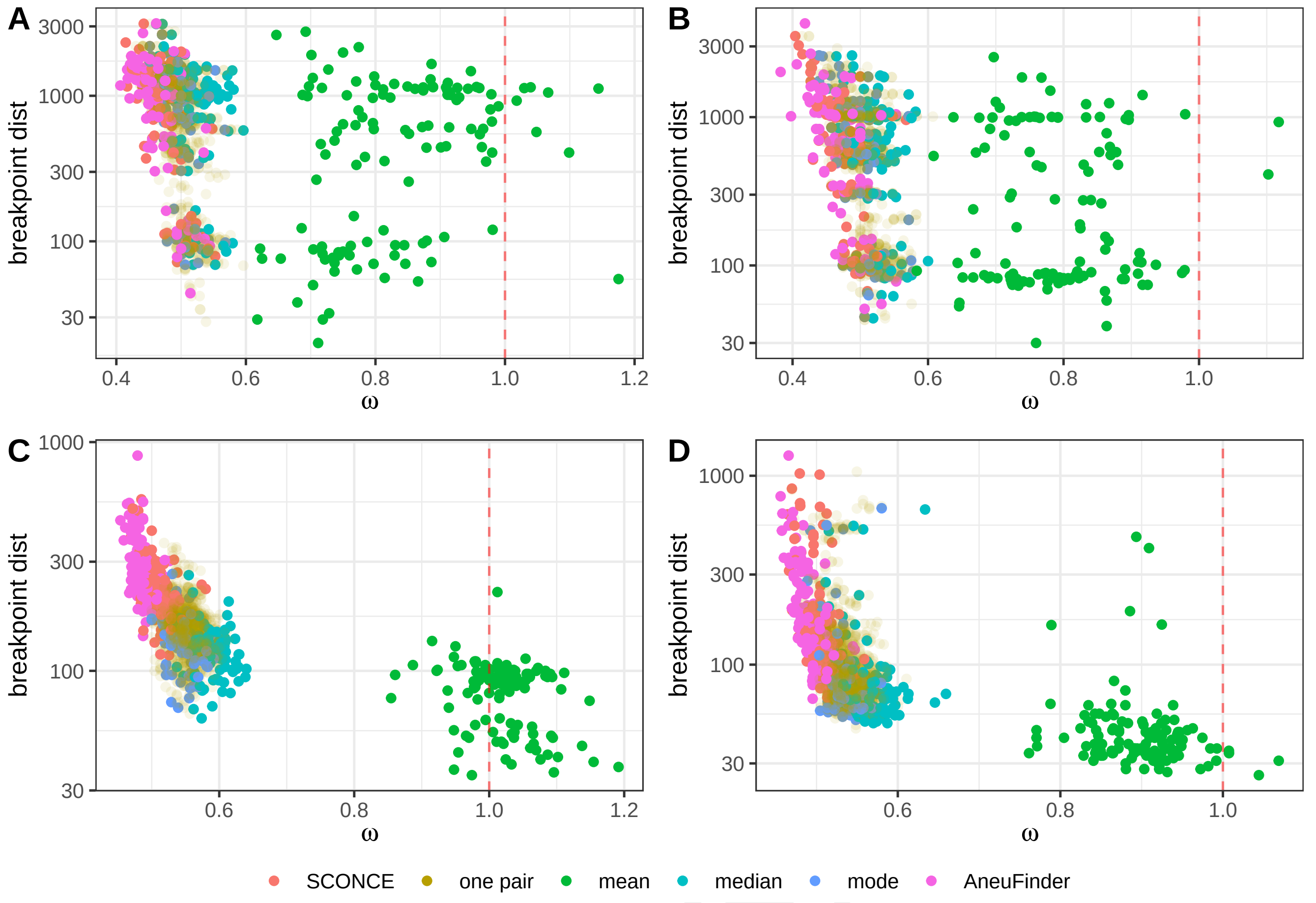


Figure S3: Breakpoint detection accuracy across parameter sets and CNP calling methods, where consensus CNPs are summarized over each cell's nearest 10 cells. Each dot represents one cell and each color represents on method, with $\omega = \frac{\#\text{ inferred breakpoints}}{\#\text{ true breakpoints}}$ on the x-axis and breakpoint distance on the y-axis. Using multiple cells consistently outperforms individual analyses.

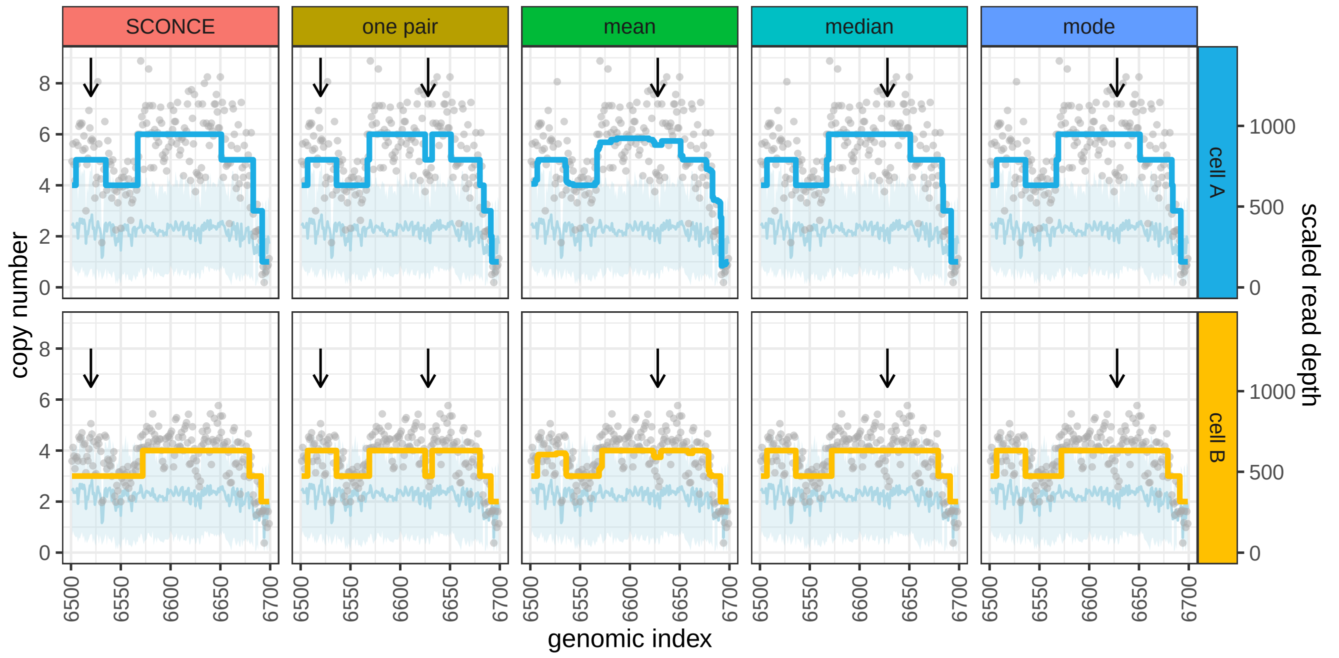


Figure S4: Improved CNA detection using multiple cells in real data (11). Copy number calls and read depths are shown across methods for two cells (cell A: SRR054596, cell B: SRR054609, from (11)) for genomic windows 6500-6700 (x-axis). Genomic windows are 250kb across hg19 and numbered sequentially. Each dot shows scaled read depth (right y-axis, relative to diploid read depth) in one window, the light blue line and band show the mean and variance of the diploid read depth data, and colored lines show inferred copy number (left y-axis). Arrows denote differences in copy number calls between methods.

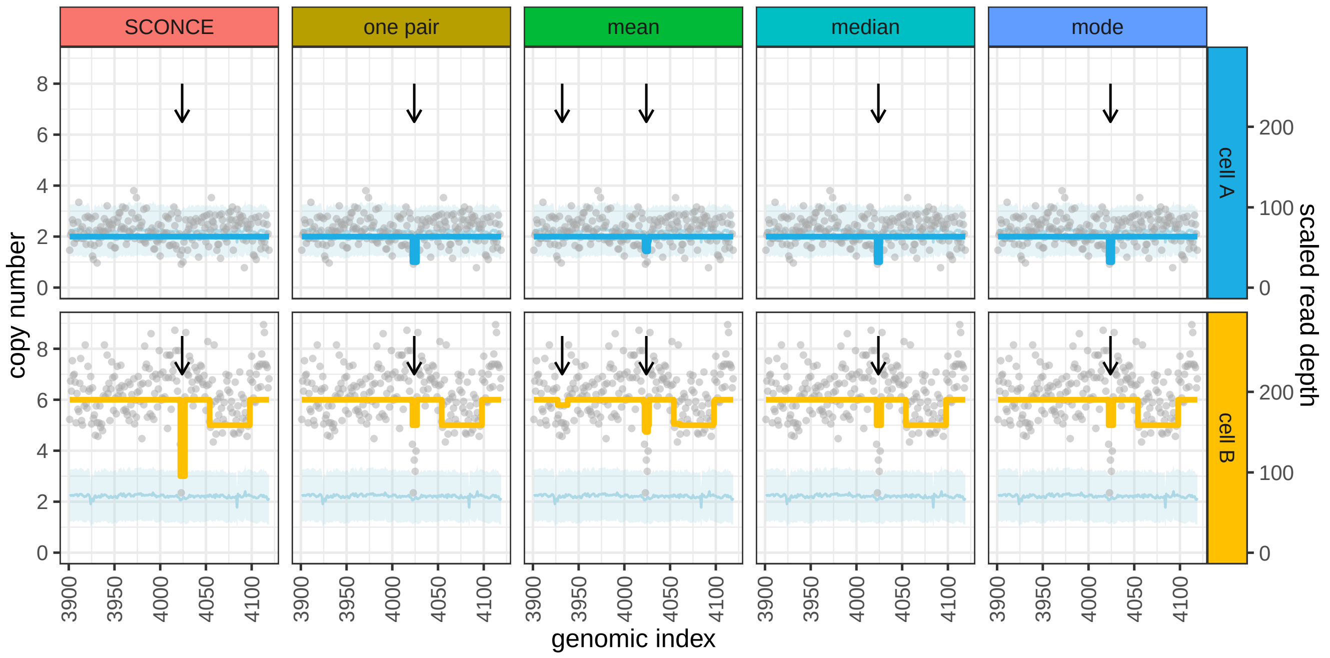


Figure S5: Improved CNA detection using multiple cells in real data (15). Copy number calls and read depth are shown for two cells (cell A: cell 1538 with barcode TAGCCGGCAAGAACTA-1 from segment D, cell B: cell 1360 with barcode GCATACAAGTAACCCT-1. from segment B, from (15)) across methods in genomic windows 3900-4120 (x-axis). Windows are numbered sequentially across hg19 and are 250kb. Each dot shows per window read depth (right y-axis, scaled to the average diploid read depth), the light blue line and band show the mean and variance of the averaged diploid read depth data, and colored lines show inferred copy number (left y-axis). Arrows denote differences in copy number calls between methods.

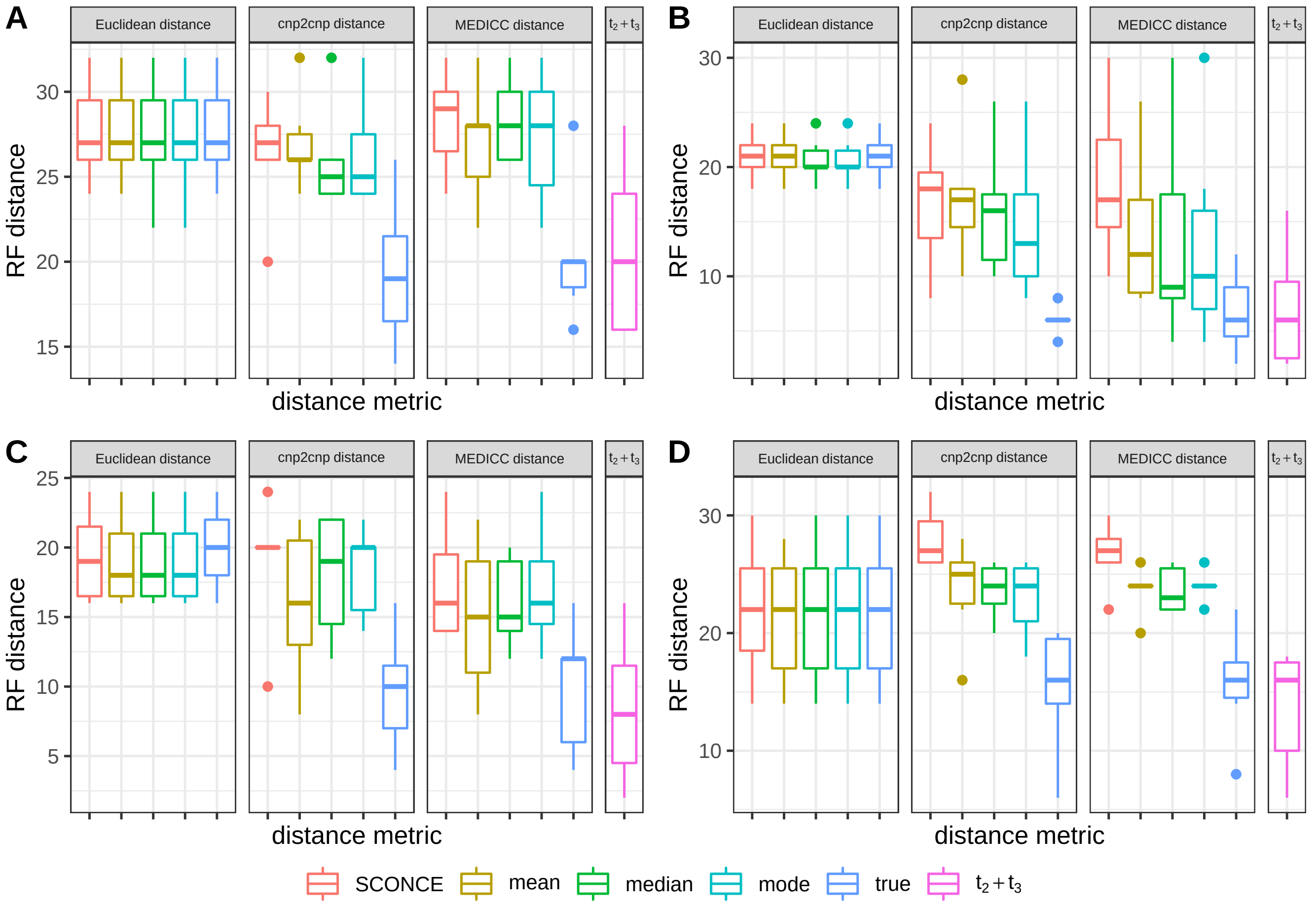


Figure S6: Robinson-Foulds (RF) distances for each simulation set (trees A-D) for different distance metrics. Distance metrics (Eucliden distance, cnp2cnp distance, MEDICC distance, $t_2 + t_3$) are grouped for each panel, and underlying CNPs sources are colored along the x-axis. Using $t_2 + t_3$ outperforms all other distance metrics on inferred copy number profiles.

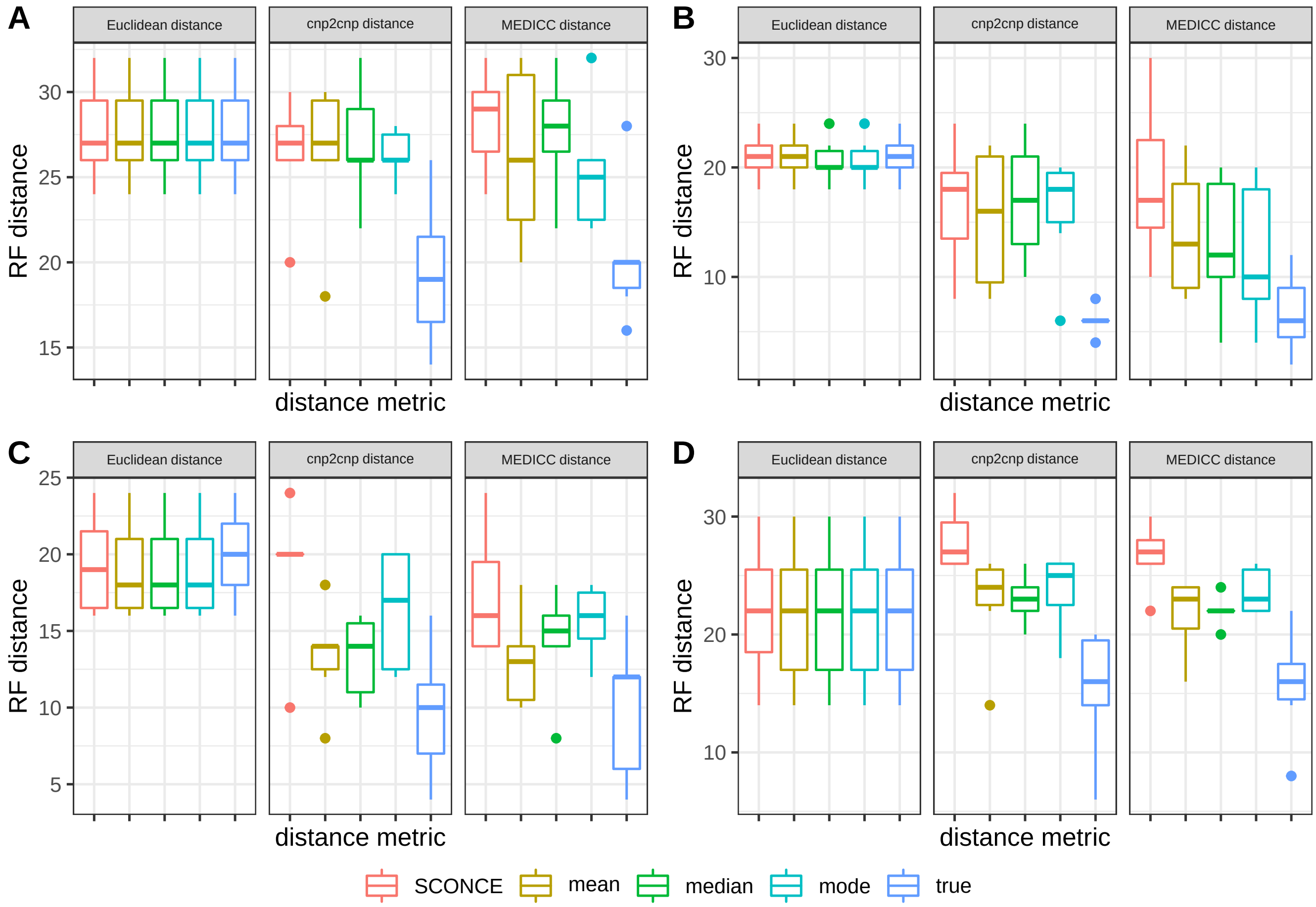


Figure S7: Robinson-Foulds distances for each simulation set (trees A-D) for different distance metrics, where consensus CNPs are summarized over each cell's 10 nearest neighbors. As in Supplementary Figure S6, distance metrics are grouped for each panel, and underlying CNPs types are colored along the x-axis. Note our $t_2 + t_3$ metric is excluded here; as a time saving measure, any pairs that aren't part of the nearest 10 of any cell (e.g., two very distant cells) are not analyzed, leading to missing data in the pairwise distance matrices.

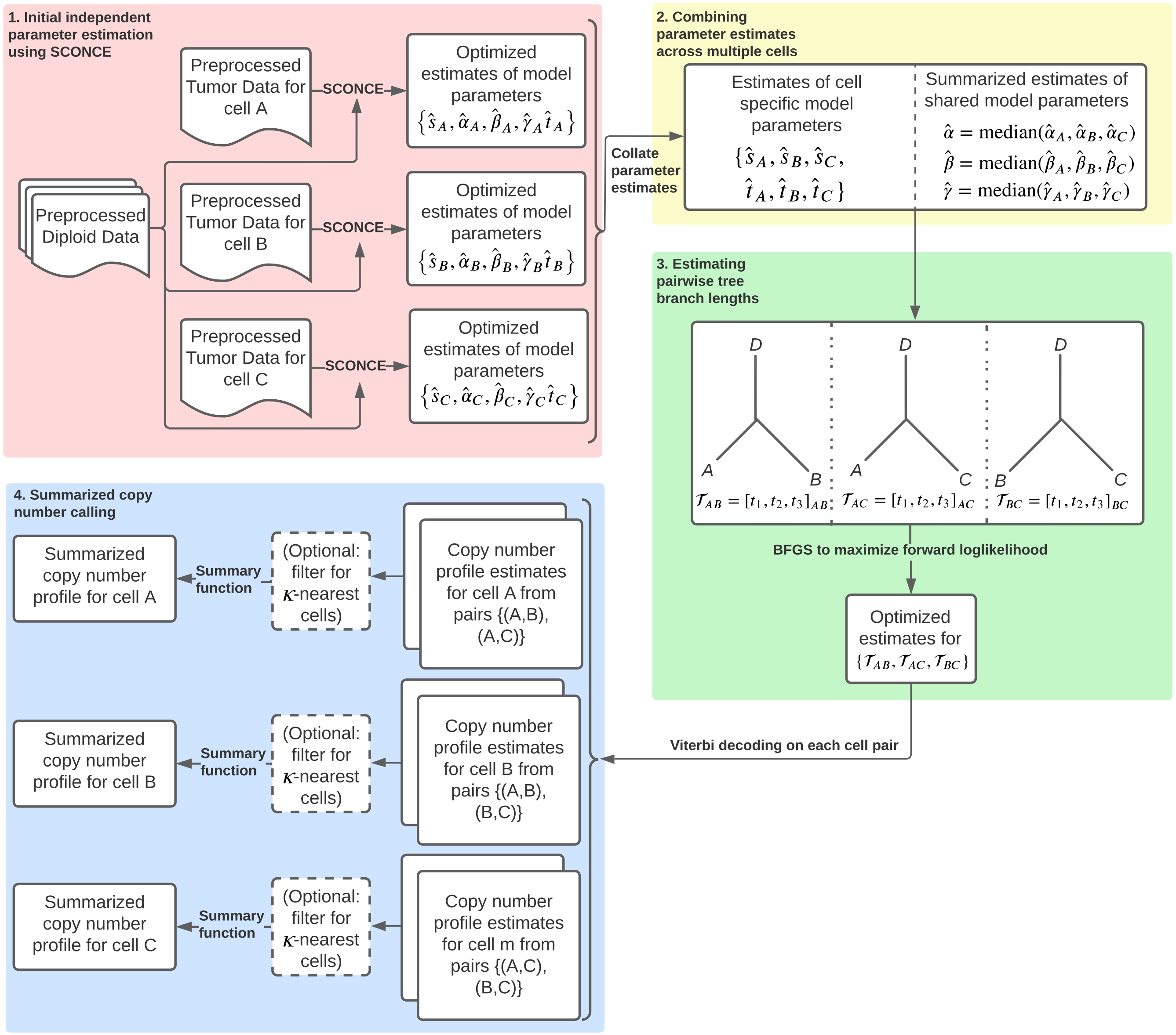


Figure S8: Detailed flowchart of the SCONCE2 pipeline. We demonstrate the pipeline with cell triplet $\{A, B, C\}$, without loss of generality. Each colored box corresponds to a step described in Detailed SCONCE2 pipeline.

### Supplementary Tables

| Simulation set/tree | Short description | Deletion rate, $\frac{\delta}{\tau_d}$ | Insertion rate, $\frac{\varphi}{\tau_a}$ | Mean deletion length, $\tau_d$ | Mean insertion length, $\tau_a$ |
|---|---|---|---|---|---|
| A | maximally imbalanced, ultrametric | 0.025 | 0.025 | 10 | 10 |
| B | perfectly balanced, ultrametric | 0.025 | 0.025 | 10 | 10 |
| C | maximally imbalanced, not ultrametric, all branches have equal length | 0.05 | 0.065 | 10.5 | 8 |
| D | maximally imbalanced, not ultrametric, terminal branch lengths decay logarithmically | 0.05 | 0.065 | 10.5 | 8 |

Table S1: Description of simulation sets under the line segment model, relative to a genome length of 100. A range of tree structures were chosen to test both typical and edge cases.

| | SCONCE | one pair | mean | median | mode | AneuFinder |
|---|---|---|---|---|---|---|
| A | 1167 | 1006 | 394 | 1018.5 | 1019.5 | 1172 |
| B | 1009.5 | 796.5 | 91.5 | 959 | 959 | 1016.5 |
| C | 206 | 153 | 84 | 142 | 142 | 263 |
| D | 153.5 | 85 | 33 | 77 | 77.5 | 168.5 |

Table S2: Full median breakpoint distance values for all trees and methods.

| | SCONCE | one pair | mean | median | mode | AneuFinder |
|---|---|---|---|---|---|---|
| A | 0.4663 | 0.49 | 0.921 | 0.4862 | 0.4856 | 0.4623 |
| B | 0.4762 | 0.5 | 0.8643 | 0.4941 | 0.4897 | 0.4744 |
| C | 0.5134 | 0.5507 | 1.1667 | 0.5467 | 0.5436 | 0.4851 |
| D | 0.504 | 0.5349 | 1.0072 | 0.5346 | 0.5338 | 0.4885 |

Table S3: Full $\omega = \frac{\#\text{ inferred breakpoints}}{\#\text{ true breakpoints}}$ values for all trees and methods.

| | SCONCE | one pair | mean | median | mode | AneuFinder |
|---|---|---|---|---|---|---|
| A | 1167 | 1014 | 591 | 1009 | 1027 | 1172 |
| B | 1009.5 | 858 | 121 | 958.5 | 959.5 | 1016.5 |
| C | 206 | 152 | 90 | 130.5 | 135 | 263 |
| D | 153.5 | 84 | 38 | 70 | 76 | 168.5 |

Table S4: Full median breakpoint distance values for all trees and methods, where consensus methods summarize over the nearest 10 cells only.

| | SCONCE | one pair | mean | median | mode | AneuFinder |
|---|---|---|---|---|---|---|
| A | 0.4663 | 0.4907 | 0.8186 | 0.5155 | 0.4929 | 0.4623 |
| B | 0.4762 | 0.5 | 0.7833 | 0.5191 | 0.4941 | 0.4744 |
| C | 0.5134 | 0.5493 | 1.02 | 0.5786 | 0.5462 | 0.4851 |
| D | 0.504 | 0.5344 | 0.9027 | 0.5639 | 0.5304 | 0.4885 |

Table S5: Full $\omega = \frac{\#\text{ inferred breakpoints}}{\#\text{ true breakpoints}}$ values for all trees and methods, where consensus methods summarize over the nearest 10 cells only.

DRAFT

| | **SCONCE** | **mean** | **median** | **mode** | **true** | $\boldsymbol{t_2 + t_3}$ |
|---|---|---|---|---|---|---|
| Euclidean distance | 27 | 27 | 27 | 27 | 27 | NA |
| cnp2cnp distance | 27 | 26 | 25 | 25 | 19 | NA |
| MEDICC distance | 29 | 28 | 28 | 28 | 20 | NA |
| $t_2 + t_3$ | NA | NA | NA | NA | NA | 20 |

(a) Tree A

| | **SCONCE** | **mean** | **median** | **mode** | **true** | $\boldsymbol{t_2 + t_3}$ |
|---|---|---|---|---|---|---|
| Euclidean distance | 21 | 21 | 20 | 20 | 21 | NA |
| cnp2cnp distance | 18 | 17 | 16 | 13 | 6 | NA |
| MEDICC distance | 17 | 12 | 9 | 10 | 6 | NA |
| $t_2 + t_3$ | NA | NA | NA | NA | NA | 6 |

(b) Tree B

| | **SCONCE** | **mean** | **median** | **mode** | **true** | $\boldsymbol{t_2 + t_3}$ |
|---|---|---|---|---|---|---|
| Euclidean distance | 19 | 18 | 18 | 18 | 20 | NA |
| cnp2cnp distance | 20 | 16 | 19 | 20 | 10 | NA |
| MEDICC distance | 16 | 15 | 15 | 16 | 12 | NA |
| $t_2 + t_3$ | NA | NA | NA | NA | NA | 8 |

(c) Tree C

| | **SCONCE** | **mean** | **median** | **mode** | **true** | $\boldsymbol{t_2 + t_3}$ |
|---|---|---|---|---|---|---|
| Euclidean distance | 22 | 22 | 22 | 22 | 22 | NA |
| cnp2cnp distance | 27 | 25 | 24 | 24 | 16 | NA |
| MEDICC distance | 27 | 24 | 23 | 24 | 16 | NA |
| $t_2 + t_3$ | NA | NA | NA | NA | NA | 16 |

(d) Tree D

Table S6: Median Robinson-Foulds distances across different distance metrics on different CNPs for each simulation set.

DRAFT

| | **SCONCE** | **mean** | **median** | **mode** | **true** |
|---|---|---|---|---|---|
| Euclidean distance | 27 | 27 | 27 | 27 | 27 |
| cnp2cnp distance | 27 | 27 | 26 | 26 | 19 |
| MEDICC distance | 29 | 26 | 28 | 25 | 20 |

(a) Tree A

| | **SCONCE** | **mean** | **median** | **mode** | **true** |
|---|---|---|---|---|---|
| Euclidean distance | 21 | 21 | 20 | 20 | 21 |
| cnp2cnp distance | 18 | 16 | 17 | 18 | 6 |
| MEDICC distance | 17 | 13 | 12 | 10 | 6 |

(b) Tree B

| | **SCONCE** | **mean** | **median** | **mode** | **true** |
|---|---|---|---|---|---|
| Euclidean distance | 19 | 18 | 18 | 18 | 20 |
| cnp2cnp distance | 20 | 14 | 14 | 17 | 10 |
| MEDICC distance | 16 | 13 | 15 | 16 | 12 |

(c) Tree C

| | **SCONCE** | **mean** | **median** | **mode** | **true** |
|---|---|---|---|---|---|
| Euclidean distance | 22 | 22 | 22 | 22 | 22 |
| cnp2cnp distance | 27 | 24 | 23 | 25 | 16 |
| MEDICC distance | 27 | 23 | 22 | 23 | 16 |

(d) Tree D

Table S7: Median Robinson-Foulds distances across different distance metrics on different CNPs for each simulation set, where consensus CNPs are summarized over only the nearest 10 cells. Note that because summarizing over only the nearest 10 cells does not guarantee all pairs of cells will be jointly analyzed, no $t_2 + t_3$ values are given here, as estimating phylogenies using $t_2 + t_3$ requires a complete distance matrix.

DRAFT
